# Interplay of action-based prediction and top-down attention: EEG evidence for joint modulation of late perceptual processing

**DOI:** 10.64898/2026.01.28.702103

**Authors:** Thomas Holstein, Jean-Christophe Sarrazin, Bruno Berberian, Andrea Desantis

**Author notes:** Co-senior authors (Bruno Berberian, Andrea Desantis). Corresponding author: Thomas Holstein.

## Abstract

Voluntary action shapes our perception through sensory predictions of its consequences. These predictions are thought to inhibit perceptual processing of predicted outcomes, leading to sensory attenuation. However, some studies have reported findings that contrast with this effect, suggesting that the influence of predictions may reflect, or interact with, attentional processes. Here, we investigated the interplay of action-based prediction and top-down attention on early and late perceptual processing, where prediction and attention refer to the likelihood of a sensory event and its behavioral relevance, respectively. Electroencephalography (EEG) was recorded while participants viewed two sequential gratings. The orientation of the first was either predicted and self-generated or unpredicted and externally generated. Participants judged the tilt direction of the second grating relative to the first but responded only when the grating appeared in a task-relevant color. EEG analyses revealed no modulation of early visual potentials (N1a), but modulations in later processing stages (P3b). Specifically, the P3b exhibited reduced amplitude for predicted and self-generated stimuli compared to unpredicted and externally generated ones, but only when they were task-relevant. Multivariate pattern analysis further showed that the significant temporal cluster supporting decoding of the first grating’s orientation was largest for relevant, predicted and self-generated stimuli. Our results suggest an optimization process whereby action-based prediction, when aligned with task goals, reduces the amount of evidence needed and increases its accumulation speed, while preserving sensory representations as accurate as in the absence of prediction/self-generation.

**Highlights:**

- Feature-based attention and prediction were orthogonally manipulated in an agency context.
- No early sensory modulation by attention or action-based prediction was observed.
- Reduction of both P3b amplitude and latency emerged for predicted/self-generated and task-relevant stimuli.
- Multivariate analysis and behavior results support an interpretation whereby relevant action-based predictions optimize evidence accumulation process.

## 1. Introduction

The ability to predict sensory information is essential for perception and sensorimotor control. Influential models of motor control propose that forward models use an efference copy of the motor command to predict the sensory consequences of voluntary actions (Crapse & Sommer, 2008; Fukutomi & Carlson, 2020; Sperry, 1950; von Holst, 1954; D. M. Wolpert et al., 1994, 1995, 1997). Interestingly, these predictions are essential not only for sensorimotor control but also contribute to the sense of agency, i.e., the subjective experience of causing one’s actions and their outcomes (Haggard, 2017; Haggard & Chambon, 2012). In fact, when sensory consequences match predicted effects, they are more likely interpreted as self-generated, whereas a mismatch suggests external causation (Barlas & Kopp, 2018; Blakemore et al., 2002; Ebert & Wegner, 2010).

Predictions generated within the motor system are also thought to impact sensory processing, in particular, the perception of time, the perceived intensity of sensory inputs and the way in which visual attention is directed (Wen & Imamizu, 2022). The phenomenon of sensory attenuation illustrates this link between prediction, voluntary action and perception. Research has shown that self-generated stimuli are typically attenuated in both subjective experience (Blakemore et al., 1999; Roussel et al., 2013; Weiss et al., 2011) and neural responses (Bäß et al., 2008; Blakemore et al., 1998; Gentsch & Schütz-Bosbach, 2011; see Hughes et al., 2013 for a review), compared to externally generated ones. This sensory attenuation is commonly interpreted as a result of action-based predictions, which allow the brain to cancel sensory responses to predicted outcomes (cf. cancellation account; Press et al., 2023), minimizing redundant information and emphasizing unpredicted events that might be more informative to update internal models (Bays & Wolpert, 2007; Brooks & Cullen, 2019).

However, research has also pointed out that the relationship between action, prediction, and sensory processing is far from straightforward (Dogge, Custers, et al., 2019; Press & Cook, 2015). For instance, some studies observed that self-generated and predicted stimuli can enhance perceptual discrimination (Desantis et al., 2014) or even are perceived with greater intensity (Kiepe et al., 2023; Thomas et al., 2022) and elicit stronger neural responses compared to externally generated stimuli or incongruent action-effects (Ackerley et al., 2012; Hughes & Waszak, 2011; Reznik et al., 2014; Simões-Franklin et al., 2011). Contrasting findings have also been observed outside the domain of action control, in studies where sensory predictions were manipulated through probabilistic tasks or external cues (see Feuerriegel et al., 2021; Press et al., 2020 for reviews).

Several accounts have been proposed to explain these discrepancies. Some suggest that, rather than being cancelled, predicted sensory inputs are pre-activated (cf. preactivation account; Roussel et al., 2013), or sharpened thereby enhancing perceptual precision (cf. sharpening account; Kok, Jehee, et al., 2012). Others propose that some effects of predictions may reflect, or interact with, attentional processes (Schröger et al., 2015; Summerfield & Egner, 2009). This latter distinction is particularly important because attention – driven by the behavioral relevance of an event – and prediction – which refers to the likelihood of that event – are often conflated (Rungratsameetaweemana & Serences, 2019; Summerfield & de Lange, 2014).

Attention is known to enhance the processing of the task-relevant stimuli by amplifying neural responses, which can enhance both the perceived intensity and perceptual discrimination (Carrasco, 2011; Desimone & Duncan, 1995; Treue, 2001). If studies focusing on sensory predictions do not adequately control for attention through behavioral relevance, observed effects on sensory processing may reflect the presence or absence of attention rather than prediction itself (Battistoni et al., 2017; Summerfield & Egner, 2009). Moreover, attentional mechanisms have often been examined using probabilistic cues (Posner, 1980), making it difficult to disentangle the effects of attention from those of prediction.

Studies that have attempted to isolate predictive mechanisms from attentional processes have reported interactive effects (Hsu et al., 2014; John-Saaltink et al., 2015; Kok, Rahnev, et al., 2012; Marzecová et al., 2017, 2018). For example, Kok et al. (2012) showed that visual neural responses to predicted stimuli were attenuated relative to unpredicted ones, particularly when visual stimuli were unattended. However, this pattern reversed when the same stimuli were attended. The authors interpreted this crossover interaction through the lenses of the predictive coding theory (Feldman & Friston, 2010; Rao & Ballard, 1999). According to this theory, attention enhances the precision (i.e., reliability) of ascending prediction error (PE) signals by boosting synaptic gain of PE units. Kok and De Lange (2015) postulated that attentional gain could be higher for PE units specifically tuned to the current prediction. As a result, when a stimulus is both predicted and attended, attention can reverse the usual prediction-suppression effect: by boosting the precision of the relevant ascending PE signals, it amplifies, rather than attenuates, the cortical representation of the predicted input (Kok, Rahnev, et al., 2012).

However, most studies examining the relationship between attention and prediction have focused on spatial attention in passive contexts. Little is known about how feature-based attention and feature-based prediction interact (Smout et al., 2020), particularly in contexts where individuals are active agents who generate predictions through voluntary actions (Jones et al., 2013). A better understanding of this interaction is mandatory to better explain the contradictory results obtained regarding the impact of voluntary action control on perception (i.e., attenuation or enhancement). The present study aims at filling this gap by combining visual psychophysics and electroencephalography (EEG) to investigate how feature-based attention and prediction shape the processing of visual stimuli generated by voluntary actions.

For this purpose, we designed an experiment that orthogonally manipulated top-down attention and action-based prediction by associating each process with a distinct stimulus feature. While EEG was recorded, participants viewed two successive gratings and reported (in some trials) whether the second was tilted clockwise or counterclockwise with respect to the first. Importantly, the orientation of the first grating was either predicted by an action (*Active* condition) or not (*Passive* condition). Top-down attention was manipulated through the color of the first grating, which determined whether the trial was relevant (i.e., required a perceptual judgment) or irrelevant (no perceptual judgment required). The relevance cue indicating which color required a perceptual judgment was provide at the beginning of each block. By assigning attention and prediction to distinct stimulus features (color and orientation, respectively), our approach allowed for a more precise investigation of their independent and interactive effects on sensory processing, compared to studies that manipulated both processes through the same feature, such as spatial location (Kok, Rahnev, et al., 2012; Marzecová et al., 2017; Wyart et al., 2012).

To investigate the effects of prediction and attention along different stages of the processing hierarchy, we analyzed early and late Event-Related Potential (ERP) components associated with these mechanisms, particularly the anterior N1 (N1a) and the parietal P3 (P3b). The N1a reflects early sensory processing and can be modulated by top-down attention (He et al., 2004, 2008; Vogel & Luck, 2000) and action-based prediction (Gentsch & Schütz-Bosbach, 2011), and can be influenced interactively by both attention and predictions (Marzecová et al., 2018). The P3b is thought to reflect the updating of higher-level internal models in response to externally-generated or unpredicted stimuli compared to self-generated or predicted ones (Bednark et al., 2015; Donchin, 1981; Polich, 2007). It has also been linked to evidence accumulation processes (Kelly & O’Connell, 2013). Within predictive processing frameworks, model updating and evidence accumulation are closely related, since unpredicted stimuli, which violate predictions, increase uncertainty and trigger the need to accumulate more evidence before committing to a decision (Zénon et al., 2019).

Based on the *crossover interaction* observed by Kok et al. (2012), we expected smaller N1a amplitudes for predicted and self-generated stimuli (*Active* condition) compared to unpredicted and externally-generated stimuli (*Passive* condition), but only when no perceptual judgment was required (*Irrelevant* condition). Conversely, in the *Relevant* condition, we expected larger N1a amplitudes in the *Active* compared to the *Passive* condition.

Regarding the P3b, we expected higher amplitudes for unpredicted events (*Passive* condition) compared to predicted stimuli (*Active* condition), reflecting greater demands for evidence accumulation or internal model updating when sensory outcomes generate larger prediction errors. Moreover, we hypothesized a *synergistic interaction* between attention and prediction, such that attention would amplify this effect, reflecting a higher priority for model updating in task-relevant trials. This hypothesis is grounded in the notion that if attention increases the gain of prediction error units (Feldman & Friston, 2010), it could lead to stronger model updating in higher-order processing areas, particularly when stimuli are task-relevant.

While ERP analyses inform about the amplitude of neural activity, they provide limited insight into the content of the information being processed. Accordingly, we performed additional exploratory analyses using Multivariate Pattern Analysis (MVPA) to examine the impact of prediction and attention on information processing. Previous studies have shown that predicted visual features (e.g., different stimulus orientations) can be decoded more accurately than unpredicted features (Kok, Jehee, et al., 2012; Yon et al., 2018), supporting the sharpening hypothesis, which suggests that prediction enhances the precision of sensory representations rather than merely cancelling sensory units encoding predicted features (cf. cancellation account; Press et al., 2023). Similar improvement in decoding accuracy has been observed for attended features compared to unattended ones (Jehee et al., 2011; Kamitani & Tong, 2005; Liu et al., 2011; Serences et al., 2009). These findings raise the question of whether top-down attention drives sensory effects attributed to prediction, interact with them, or operates independently (Smout et al., 2019). Based on the synergistic interpretation of the interaction between attention and prediction, we hypothesized that prediction would improve the decoding of stimulus orientation by sharpening sensory representations, while attention would further enhance decoding by increasing the gain of task-relevant neural populations.

## 2. Materials and methods

### 2.1. Participants

A sample size of twenty-four participants was estimated to achieve a statistical power of ∼0.80, with the significance level of p = 0.05. This calculation was based on studies by Marzecová et al. (2017, 2018), who reported medium-to-large effect sizes when examining the impact of prediction and task relevance on N1 and P3 components using a design similar to ours. This number also correspond to the sample typically used in previous experiments investigating sensory attenuation of visual events (cf. Hughes & Waszak, 2011 with 18 observers; Gentsch et al., 2012 with 23 observers). Twenty-five volunteers took part in the experiment in return for a gift card worth 15€ per hour of experiment. One participant was excluded from the analyses due to discrimination performance being too close to the chance level, falling outside our inclusion criteria (average discrimination performance should be between 60-90% for each block). This left a final sample of twenty-four observers (10 female, total mean age: M=30.3 SD=8.3). All participants had normal or corrected-to-normal visual acuity, declared no oculomotor, neurological or psychiatric disorders and were naïve as to the hypotheses under investigation. The experimental session lasted between 2 hours and 2.5 hours, including the training, the staircase procedure, the setting up of the EEG, the main session, and breaks. Participants gave written and informed consent before participating. The study was approved by the Ethics Committee of Université Paris Cité (IORG0010120-2025-77) and conducted in agreement with the requirements of the Declaration of Helsinki.

### 2.2. Materials and stimuli

Participants sat in front of a 19-inch cathode-ray tube (CRT) monitor (resolution of 1024 x 768 pixels refresh rate of 60 Hz). Head movements were restricted by a chin rest located 70 cm away from the screen. The experiment was programmed with Python (Python Software Foundation, Python Language Reference, version 3.11) and stimulus presentation was controlled by Psychopy 2023.1.3 (Peirce et al., 2019).

The stimulus consisted of a sinusoidal grating with a circular luminance profile (spatial frequency of 2.0 cycles/degree of visual angle (dva), fixed phase of 0, size of 7 dva) and a circular noise patch (size of 7 dva)). The noise patch consisted of a pixel matrix containing values between 0 and 1 and following a folded-normal distribution with a mean of 1 and a standard deviation of 0.4. Value of 1 corresponding to the color of the grating (red or blue) and 0 to the grey background color (RGB: 0.5, 0.5, 0.5). This distribution was chosen in order to enhance color discriminability. In line with this strategy, the two colors were set to have the most separated chromaticity values while keeping similar luminance and contrast values with the background color. Therefore, the red grating (RGB: 0.6, 0, 0) and the blue grating (RGB: 0,0,1) had an opacity of 0.8 and 0.85 respectively. The noise patch was displayed in front of the grating with a contrast and opacity of 60% (100% being fully opaque and 0 fully transparent). Using a Colorimeter (Cambridge Research Systems ColorCAL MKII) the red and blue stimuli (grating + noise) were calibrated to have matched luminance, each measuring approximately 3.94 cd/m². A white fixation point of 1 dva was displayed at the center of the screen and of the stimulus.

While participants were performing the behavioural task, continuous EEG signals were recorded using 64 Ag/AgCl electrodes mounted on an elastic cap and amplified by an ActiCHamp Plus amplifier (Brain Products GmbH, Munich, Germany). Electrodes were arranged according to the international 10–20 systems. EEG data were record without reference and were offline referenced to the average. Four electrodes were deported to record EOG signals: Fp1 and Fp2 were placed above and below the right eye, respectively, to record vertical saccades and blinks; while AF7 and AF8 were placed on both temples to measure horizontal saccades. EEG signals were sampled at 500 Hz.

### 2.3. Procedure

#### 2.3.1. Main task

Participants performed an equiprobable go/no-go task within a two-interval forced choice (2IFC) discrimination task. The main task consisted of 10 blocks of 56 pseudo-randomized trials each. Each trial involved the presentation of two successive gratings, displayed at the center of the screen for 50 ms each, and separated by a time interval of 800 ms from the end of the first grating (Figure 1). The first grating was oriented at either 45° (right) or -45° (left) relative to the vertical line. The second grating was slightly tilted clockwise or anticlockwise with respect to the first. The orientation difference was determined individually for each participant during a preliminary phase to ensure a performance level of approximately 75% correct responses (see 2.3.3. Staircase Procedure).

**Figure 1.**
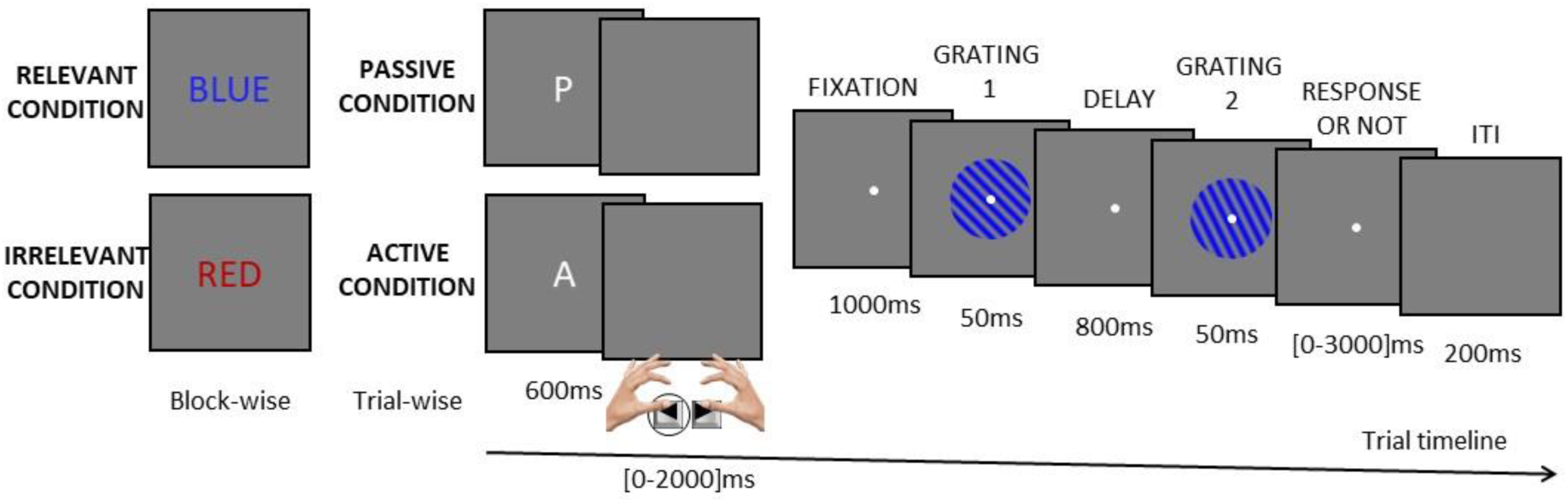
Schematic representation of the task showing both active and passive conditions. In the active trial illustrated here, participants generated the first grating oriented -45° by pressing the left button. In this example, the blue gratings indicate that the trial is relevant. The relevance cue was presented at the beginning of each block.

In *Relevant* trials participants were required to report whether the second grating was tilted clockwise or anticlockwise with respect to the first by pressing designated left or right keys, respectively, with their index fingers. Instead, in *Irrelevant* trials, they were not required to provide any explicit judgments, but were still asked to observe the presented stimuli. No feedback on discrimination performance was provided during this phase. However, an error message was displayed if participants responded in irrelevant trials or failed to respond in relevant trials.

The color of the grating indicated whether participants were completing an irrelevant or a relevant trial. Both gratings within a trial had the same color, which could be either red or blue and varied from trial to trial. The relevant color was indicated at the start of each block and changed between blocks.

Importantly, the orientation of the first grating could be either predictable (*Active* condition) or unpredictable (*Passive* condition). Specifically, in the active condition, participants determined the orientation by pressing a button: pressing the left button with their left thumb generated a -45° tilt, while pressing with their right thumb the right button generated a 45° tilt. Participants were instructed to execute left and right actions approximately equally often throughout the task, and they received feedback at the end of each block about the proportion of left and right actions performed. In the passive condition participants performed no action, the orientation of the first grating was determined by the computer, with the two orientations being equiprobable throughout the task.

The *Active Relevant*, *Passive Relevant*, *Active Irrelevant* and *Passive Irrelevant* trials were pseudo-randomized such that each condition was equally represented within each block (25% of the trials per condition), while their order of presentation was randomized. Participants were cued whether they had to complete a passive or an active trial by the presentation of the letter “A” or “P”, respectively. This cue remained visible for 600 ms. In active trials, after the cue disappeared a blank screen was presented. Participants had a maximum of 2000 ms to perform their action. Once the action was performed, the white fixation point reappeared, and the grating was displayed 1000 ms later. If no action was performed within the time limit, the trial was aborted and an error message was displayed. The aborted trial was replayed later in a random order within the same block. In passive trials, the onset of the fixation point after the blank screen was adjusted on a trial-by-trial basis to match action latency of the active trials, ensuring similar temporal relations dynamics across the two conditions.

#### 2.3.2. Training Phases

Before the main task, participants performed trainings to familiarize themselves with the task structure and then agency conditions. This training consisted of two phases:

- **Initial Training:** Participants completed 2 blocks of 16 pseudo-randomized trials each, for a total of 32 trials. The trials included equal occurrences of all stimulus orientations, rotations, and colors. In this phase, the orientation of the second grating was fixed at 10° clockwise or anticlockwise relative to the first grating. Participants received visual feedback on their responses, indicating correct or incorrect discrimination in relevant trials and correct or incorrect absence of responses in irrelevant trials. All trials were conducted in the passive condition, with no agency over the orientation of the first stimulus.
- **Second Training:** Participants completed 4 blocks of 8 pseudo-randomized trials each, totaling 32 trials. The first 2 blocks used a 10° orientation difference, while the last 2 blocks used individual thresholds derived from the staircase procedure. During this training, we introduced the active condition.

#### 2.3.3. Staircase Procedure

The orientation difference between the two gratings was adapted for each participant through a staircase procedure to determine their discrimination threshold. The task during this phase was identical to the one performed in the passive trials of the main task, where participants could not predict the orientation of the first grating. To ensure that individual discrimination thresholds were obtained under conditions as similar as possible to the main task, both relevant and irrelevant trials were reproduced during this phase.

Two interleaved adaptive staircases using the QUEST algorithm (Watson & Pelli, 1983) controlled the orientation difference between the two gratings: one starting with a high orientation difference (5°), the other with a low orientation difference (0.5°). This procedure was conducted separately for each grating color (red and blue), with the order counterbalanced across participants.

The orientation difference was adjusted based on participants’ responses, aiming to converge on a threshold corresponding to ∼75% correct performance. Each staircase stopped after 50 trials or when the orientation threshold converged with a confidence interval of 85%.

### 2.4. EEG analysis

#### 2.4.1 EEG-preprocessing

EEG pre-processing was performed with Python (Python Software Foundation, Python Language Reference, version 3.11) and the MNE toolbox (Gramfort, 2013). Based on recent recommendations for ERP analyses, EEG signals were filtered using a non-causal Butterworth filter with a 30 Hz low-pass cutoff and a 0.1 Hz high-pass cutoff frequencies, applying a slope of 12 dB/octave (Zhang et al., 2024). Noisy channels were replaced by an interpolated weighted average from surrounding electrodes. Zero to 3 channels were interpolated per participant with an average of 0.4 channel which appeared to no longer record any physiological signal (cf. Supplementary Material). Data were re-referenced to the average reference. Independent Component Analysis (ICA) was used to remove eye-blinks and saccades from continuous data, based on correlation with EOG channels and on visual inspections of topographic maps, power spectrum, reconstructed latent sources and the EEG data reconstruction.

Epochs ranging from −200 to 800 ms were time-locked to the onset of the first grating. Baseline correction was applied using the 200 ms window preceding stimulus presentation. Artifact rejection was conducted through visual inspection with the help of an open-source automatic rejection algorithm available on Python that estimates sensor-specific peak-to-peak thresholds by Bayesian optimization and cross-validation (Jas et al., 2017). This led to the removal of an average of 3.84% (SD: 3.93%), 3.07% (SD: 3.37%), 3.9% (SD: 3.41%) and 4.29% (SD: 4.01%) for the *Active Relevant*, *Passive Relevant*, *Active Irrelevant* and *Passive Irrelevant* conditions, respectively.

#### 2.4.2. ERP analyses

We report the results for the N1a and P3b components observed after the onset of the first stimulus. Other components, such as the P1, N170, P2, N2 and P3a were also analyzed but reported in the Supplementary Material, as they did not yield conclusive results. For each component, we identified the maximum peak amplitude for each participant and condition within a specific time window defined based on literature (Luck, 2014), across all electrodes of interest. Visual inspection was conducted to ensure that the component peaks were contained within their respective time windows for each participant and electrode of interest.

The peak of the N1a component was identified within a 100-200 ms post-stimulus time window. Similar to the study by Marzecova et al. (2018), we have conducted statistical analyses on a cluster of six fronto-central electrodes (F1, F2, Fz, FC1, FC2, FCz). We also conducted statistical analyses on the electrode where the maximum negative peak was reached in the grand average, i.e., on AFz electrode. Statistical analyses for the P3b were conducted at Pz electrode, where the maximum positive peak was observed and which is the classical electrode used for analyzing the P3b. The peak of the P3b was identified within a time window of 350-500 ms post-stimulus. In addition to the amplitude analyses, we also performed an exploratory analysis of P3b latency, defined as the time point of the maximum positive peak within the P3b time window for each participant.

Normality of amplitude distributions was assessed using the Shapiro–Wilk test (Shapiro & Wilk, 1965). For the N1a component, which violated normality assumptions, a Yeo–Johnson transformation was applied across all conditions (Yeo & Johnson, 2000). Statistical analysis was conducted using repeated measures ANOVAs on peak amplitudes at each electrode and electrode cluster of interest. The within-subject factors included task-relevance (relevant, irrelevant) and agency (active, passive). The assumptions of normally distributed and homoscedastic residuals in the repeated-measures ANOVA model have been confirmed through visual inspection of QQ-plots (Quinn & Keough, 2002) and by applying Shapiro–Wilk tests to the model residuals. In cases where the ANOVA revealed significant interactions, pairwise t-tests were conducted, with Bonferroni correction for multiple comparisons.

#### 2.4.3. Multivariate Pattern Analysis (MVPA)

MVPA was conducted on the same pre-processed EEG data used for ERP analysis. A Linear Discriminant Analysis (LDA) classifier was applied to decode orientation of the first stimulus (left vs. right) for each condition and participant. The analysis followed a decoding-over-time approach, where the classifier was trained and tested at each timepoint using non-overlapping time windows of 50 ms. EEG signals within each window were averaged to increase signal-to-noise ratio (Huang et al., 2021). Non-overlapping windows were chosen to avoid redundancy and provide independent time points to the classifier. Additional time windows of 20 ms and 100 ms were also tested, and these results are reported in the Supplementary Material. To ensure robust and unbiased performance estimates, a 10-fold cross-validation procedure was employed. Classifier performance was quantified using the Receiver Operating Characteristic Area Under the Curve (ROC-AUC) score, averaged across folds. To evaluate whether decoding performance exceeded chance level (0.5), statistical significance was assessed using a nonparametric cluster-based permutation test with 10,000 iterations (Maris & Oostenveld, 2007).

#### 2.4.4. MVPA feature selection

Since EEG signals are complex and collected using multiple electrodes, they often contain irrelevant information for the classification (Kabir et al., 2023). To enhance decoding accuracy, feature dimensionality was reduced before the MVPA by selecting the most informative electrodes. Specifically, for each condition and each electrode, we computed the grand-average ERP for the left- and right-oriented gratings, separately. We then calculated the difference between these two averages (left and right), separately for each condition and electrode. Within each predefined time window, we computed the mean of the difference across time points to keep one single value per time window, again separately for each condition and electrode. For each time window and condition, we selected the 10 electrodes with the largest mean difference. Electrodes selected across windows were pooled together, resulting in approximately 30 electrodes per condition (Fig. 2). MVPA was performed on this pooled set of electrodes separately for each condition. Additionally, we tested alternative selections of five and fifteen electrodes per time window, as well as a classifier working on all channels. We report the corresponding results in the Supplementary Material.

**Figure 2.**
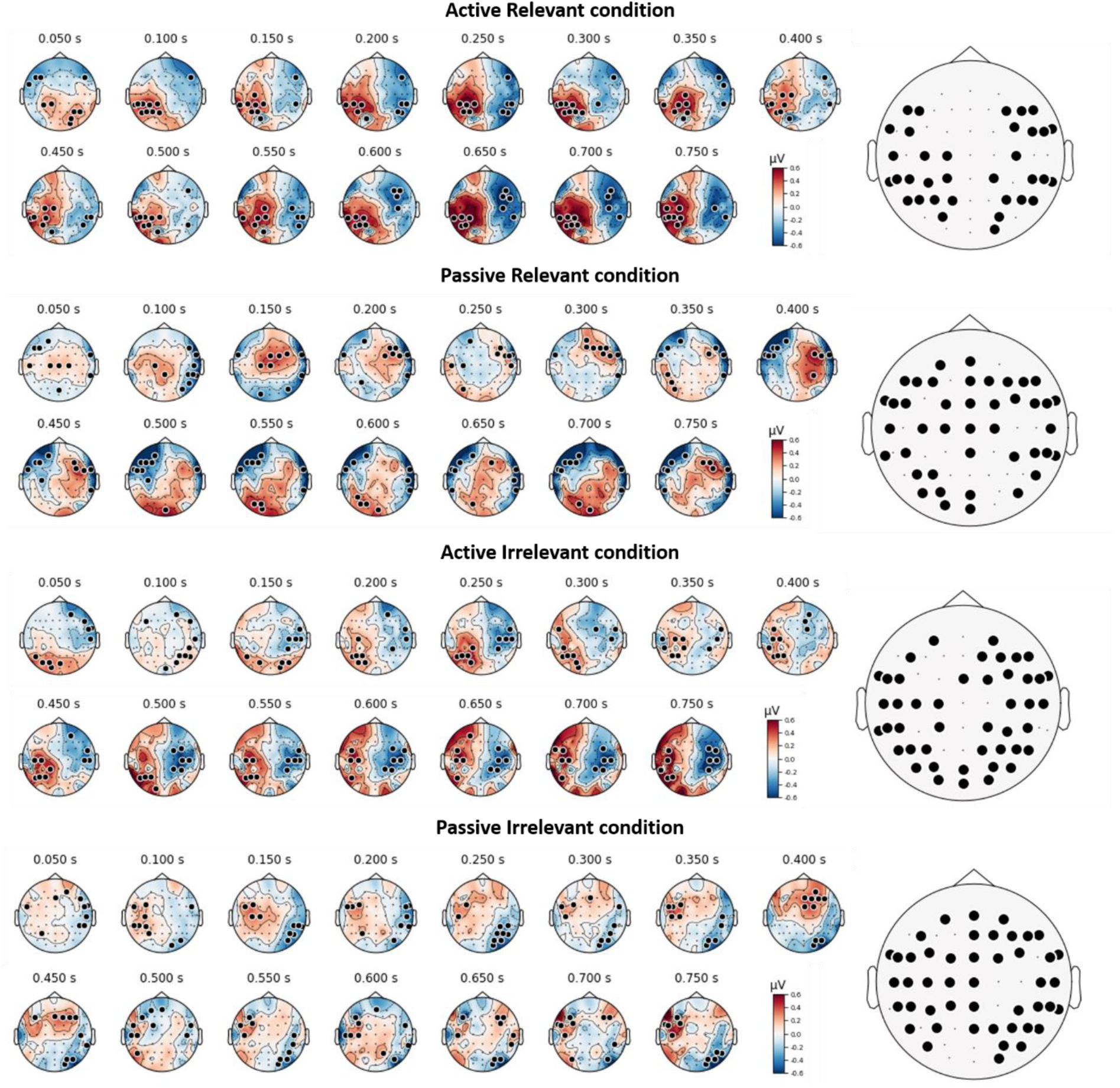
Left: Topographical maps showing the ten electrodes with the largest mean difference in evoked potentials between left- and right-oriented stimuli for each time window, separately for each condition. Electrodes were selected based on grand-average differences across participants. Right: Electrodes pooled across all time windows used for MVPA analysis, shown separately for each condition.

**Figure 3.**
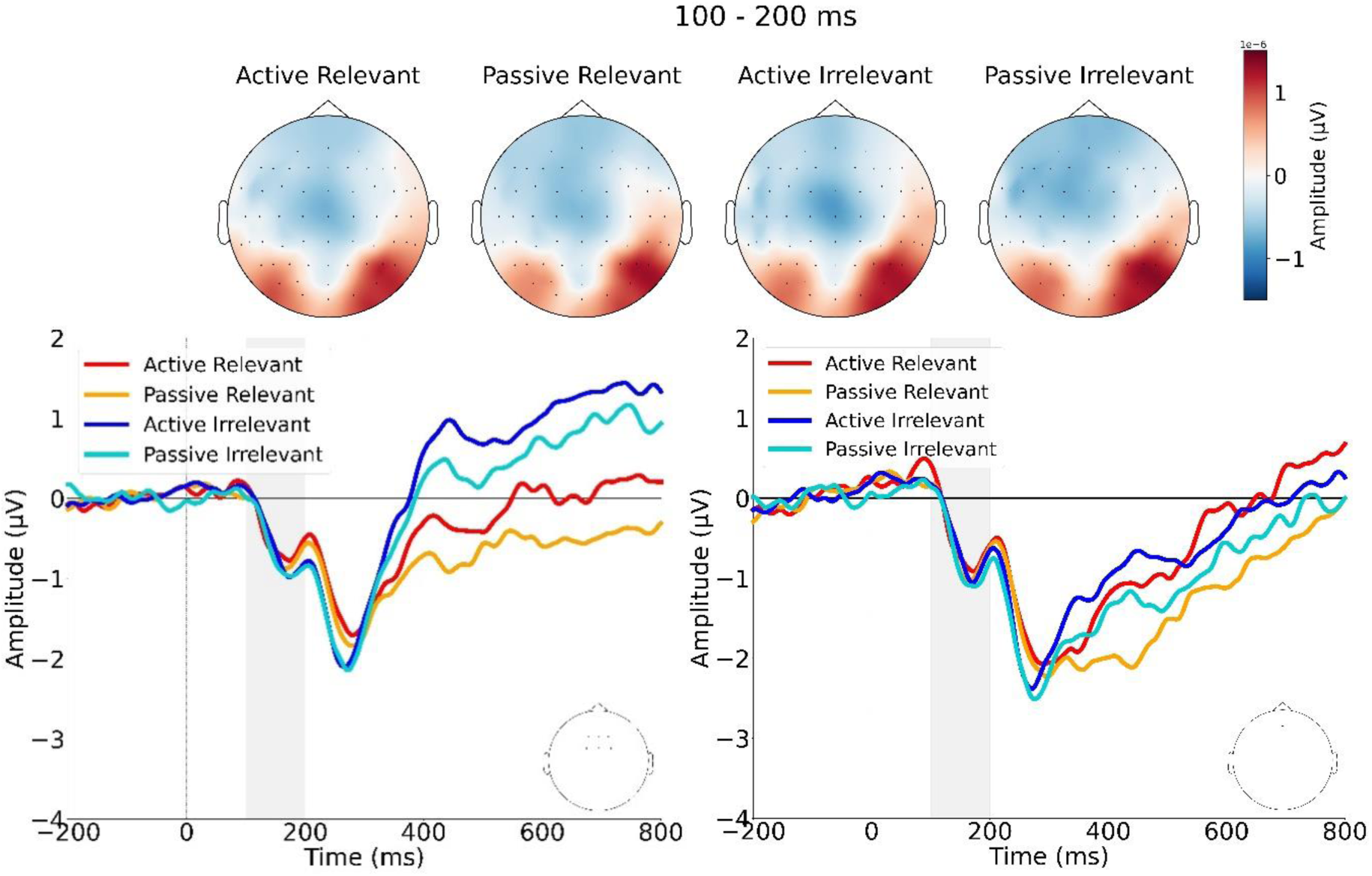
(top) Topography of brain activity averaged over the N1a component time window (100–200 ms). (left) Grand-average ERPs from the fronto-central cluster (F1, F2, Fz, FC1, FC2, FCz), (right) and from the AFz electrode, showing the N1a component across conditions. The grey areas indicate the time window (i.e., from 100 to 200 ms) used to identify the maximum negative peak for each participant and condition.

**Figure 4.**
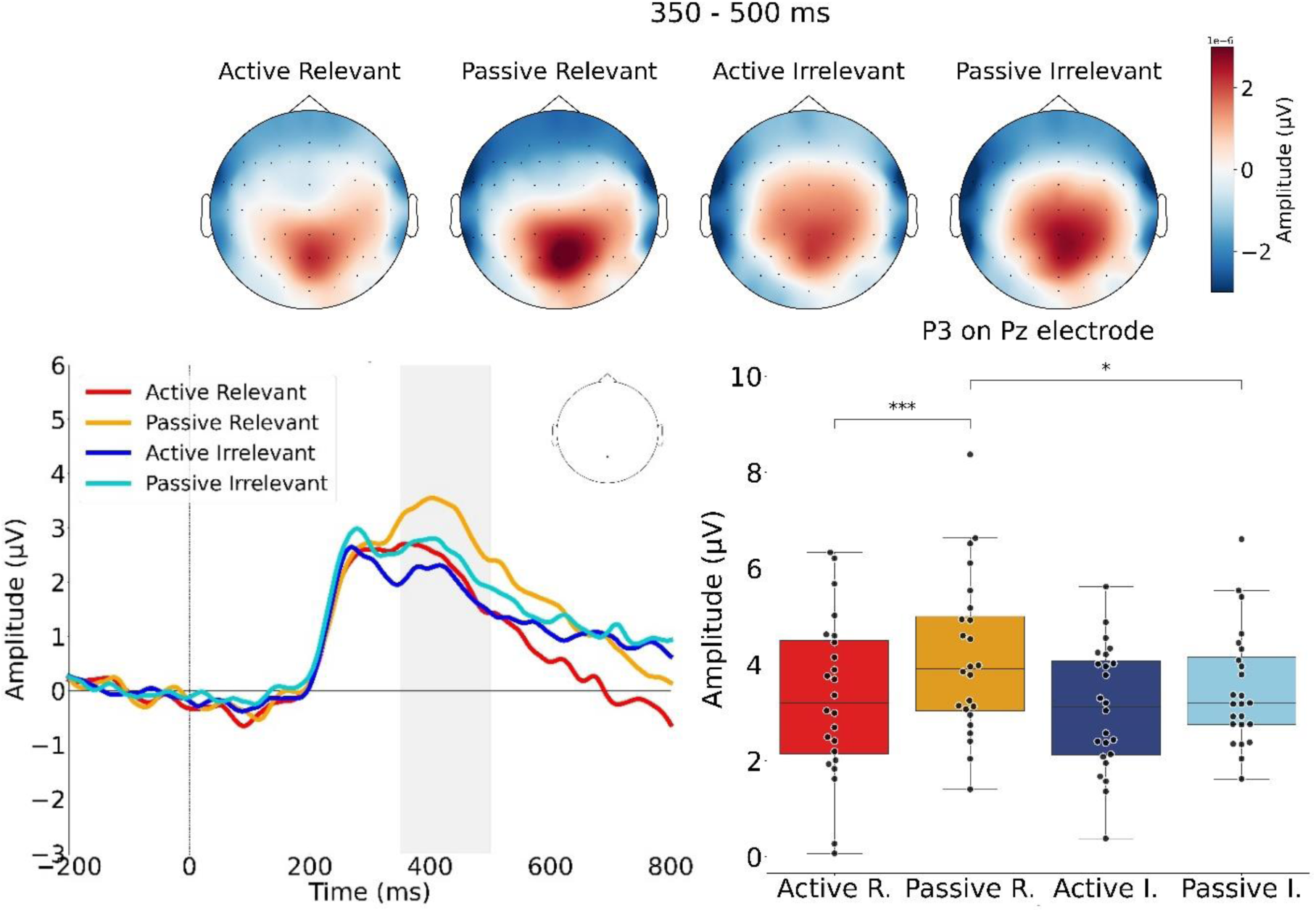
(top) Topography of brain activity averaged over the P3b component time window (350–500 ms). (left) Grand average ERPs of the P3b and (right) boxplots of P3b peak amplitude for each condition illustrating the interaction between agency and task-relevance. Error bars in the boxplots represent the interquartile range (IQR), with the central line indicating the median and the asterisks indicating levels of statistical significance after Bonferroni correction (*: .01 < p < .05; ***: p < .0001).

**Figure 5.**
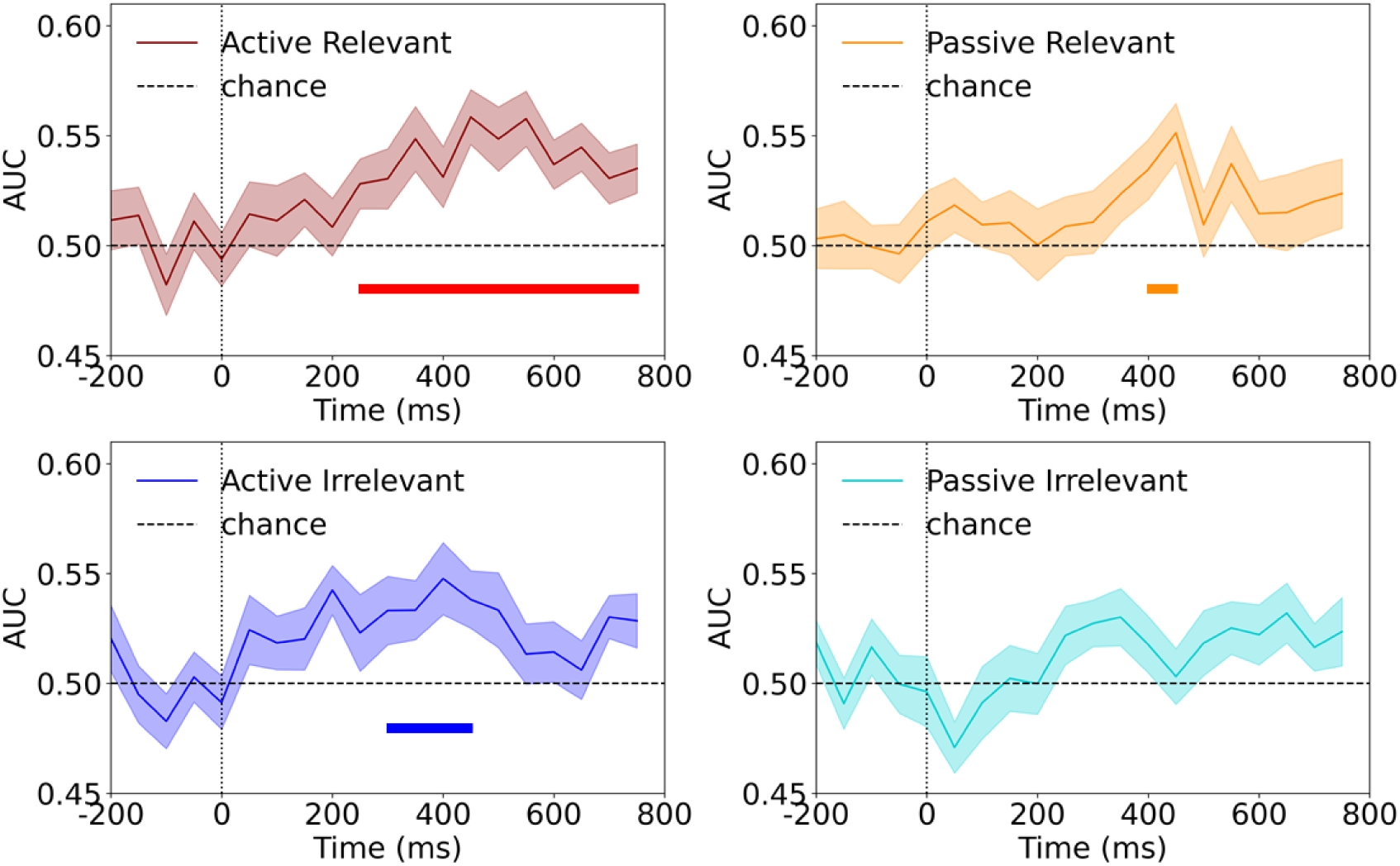
AUC scores over time for each condition (Active Relevant, Passive Relevant, Active Irrelevant, Passive Irrelevant) for the 50 ms time window classifier. Time 0 corresponds to the onset of the first grating. The shaded area around each curve represents the standard error of the mean (SEM), reflecting variability across participants. The horizontal line under the curve indicates time intervals where classification accuracy was significantly above chance (p < 0.05).

### 2.5. Behavioral data analysis

To obtain a more direct measure of task performance, we analyzed both accuracy rates and reaction times (RT). All behavioral data were normally distributed, as confirmed by Shapiro–Wilk tests, and were therefore analyzed using paired t-tests. The significance level adopted for all statistical tests was p < .05.

#### 2.5.1. Hierarchical Drift Diffusion Model

Given that accuracy remained stable across conditions while reaction times (RTs) were significantly modulated by action-based prediction, we analyzed the possibility of a speed-accuracy trade-off. To explore this, we modelled reaction times and accuracy via a Hierarchical (Bayesian) Sequential Sampling Model (HSSM) library in python (Fengler et al., in prep.). This allowed us to investigate whether evidence accumulation was more efficient in the active compared to the passive condition (cf. Ratcliff & McKoon, 2008, for a review on sequential sampling models). Specifically, we estimated four parameters: drift rate *v* (the speed of evidence accumulation), boundary separation *a* (the amount of evidence needed to reach a decision threshold), starting point *z* (the response bias) and non-decision time *t*. Each parameter was modeled as a function of agency condition (*Active Relevant* vs *Passive Relevant*) with participant-level random intercepts and random slopes for agency condition, corresponding to the following model structure: *y ∼ agency_condition + (1 + agency_condition | participant)*. Further details regarding model specification, parameterization and model diagnostics are provided in the Supplementary Materials. Posterior distributions were summarized with mean differences, 95% highest density intervals (HDI), and the proportion of posterior mass on one side of zero (P>0) for each parameter. Following recent recommendations for Bayesian reporting (Kruschke, 2021), we did not apply a binary significance threshold. Effects were classified as strong evidence when ≥ 95% of posterior mass lay on one side of zero and the 95% HDI excluded zero; weak evidence for 90 – 95% of posterior mass and very weak for 85 – 90%, even when the HDI included zero. Posterior mass < 85% was considered insufficient evidence for an effect.

### 2.6. EEG-behavior correlation analysis

To examine the correlation between electrophysiological and behavioral effects of action-based prediction, we computed the *Active Relevant* – *Passive Relevant* difference for P3b amplitude, P3b latency and reaction time (RT) for each participant. Separate linear regression models were fitted for each P3b measure, with either the P3b amplitude difference or the P3b latency difference as the dependent variable and the RT difference as the predictor. Data normality and residual normality were assessed using Shapiro–Wilk tests, and no significant deviation from normality was observed (amplitude: p = .45; latency: p = .15; RT: p = .30; residual model with amplitude: p = .65; residual model with latency: p = .16). Homoscedasticity was confirmed using the Breusch–Pagan test (p = .99 for amplitude model; p = .90 for latency model).

## 3. Results

### 3.1. Behavioral results

#### 3.1.1. Relevant action-based prediction reduces reaction time

As the data follow a normal distribution and are homoscedastic, a paired t-test was performed to compare the percentage of correct responses between *Active Relevant* (mean = 80.30, SD = 7.60) and *Passive Relevant* (mean = 79.52, SD = 7.47) conditions. No significant difference was observed (t(23) = 0.694, p = 0.494). However, a paired t-test on RT data revealed a significant difference (t(23) = -2.188, p = 0.039) with a shorter mean RT for the *Active Relevant* condition (mean = 0.693 s, SD = 0.103) compared to the *Passive Relevant* condition (mean = 0.712 s, SD = 0.104), suggesting that grating orientation might have been encoded more efficiently in the active than in the passive condition. HSSM explored this possibility and provided suggestive evidence for a higher drift rate (*v*), reflecting faster evidence accumulation, in the *Active Relevant* compared to the *Passive Relevant* condition, with 92.3% of the posterior distribution supporting this difference (mean difference = 0.093; 95% HDI = [−0.0373, 0.2209]). The boundary separation parameter (*a*) showed that 86.9% of the posterior distribution favored a smaller threshold in the *Active Relevant* compared to the *Passive Relevant* condition (mean difference = −0.012; 95% HDI = [−0.033, 0.0086]; P>0 = 0.102), suggesting very weak evidence for this difference. Non-decision time (*t*) (mean difference = − 0.0026; 95% HDI = −0.0196, 0.0142]; P>0 = 0.389) and starting point (*z*) (mean difference = −0.0054; 95% HDI = [−0.0270, 0.0175]; P>0 = 0.318) showed no credible differences. These results indicate that relevant action-based prediction does not enhance accuracy but reduces reaction time, which is mostly driven by the higher speed of evidence accumulation and the reduced amount of evidence accumulation needed to reach a decision threshold.

### 3.2. ERP results

#### 3.2.1. N1a: no action-based prediction or top-down attention modulations

A repeated measure ANOVA (agency: active vs. passive; task-relevance: relevant vs. irrelevant) on N1a calculated in the fronto-central cluster (F1, F2, Fz, FC1, FC2, FCz), revealed no significant main effect of agency (F(1,23)=0.20, p=.656) or task-relevance (F(1,23)=0.22, p=.645), and no significant interaction (F(1,23)=0.00, p=.947). Similar results were found for N1a at AFz electrode, with no significant main effect of agency (F(1,23)=0.08, p=.775), or task-relevance (F(1,23)=0.55, p=.467) and no significant interaction (F(1,23)=0.00, p=.995). Further results provided in the Supplementary Materials, specifically on P1 component, led to similar results with a null effect. These results seem to indicate that both action-based prediction and top-down attention, as manipulated here, have no impact on early sensory processing.

#### 3.2.2. P3b: prediction-suppression effect for relevant stimuli and attentional gain effect for unpredicted/externally-generated stimuli

A repeated measure ANOVA on the P3b at Pz showed a significant main effect of agency (F(1,23) = 13.51, p = .001, η^2^*_G_* = 0.045), with higher amplitude for the passive (M = 3.82 µV, SD = 1.36) compared to the active condition (M = 3.20 µV, SD = 1.39). A main effect of task-relevance (F(1,23) = 5.24, p = .032, η^2^*_G_* = .022) was also observed with higher amplitude for the relevant condition (M = 3.72 µV, SD = 1.62) compared to the irrelevant condition (M = 3.29 µV, SD = 1.13). The interaction between task-relevance and agency was also significant (F(1,23) = 6.55, p = .018, η^2^*_G_* = 0.006). Post-hoc pairwise t-tests with Bonferroni correction identified a significant difference between the *Active Relevant* condition and the *Passive Relevant* condition (t(23) = -5.34, p =.0001, d = -1.090). The P3b peak amplitude was the largest for the *Passive Relevant* condition (M = 4.15 µV, SD=1.65), following by the *Passive Irrelevant* condition (M = 3.50 µV, SD = 1.20), *Active Relevant* condition (M = 3.30 µV, SD = 1.67), and then the *Active Irrelevant* condition (M = 3.09 µV, SD = 1.29). A significant difference between the *Passive Relevant* condition and the *Passive Irrelevant* condition was also observed (t(23) = 3.26, p = .0206, d = 0.666). Regarding P3b latency, the ANOVA revealed no main effects of agency (F(1,23) = 1.86, p = .186) or task-relevance (F(1,23) = 1.19, p = .287), but a significant interaction (F(1,23) = 12.61, p = .002, η^2^*_G_* = .029). Post-hoc analyses showed shorter latencies in the *Active Relevant* condition compared to both the *Passive Relevant* (t(23) = -2.46, p = .0435, d = -0.503) and *Active Irrelevant* conditions (t(23) = -2.49, p = .0435, d = -0.508). No other comparisons indicated significant difference. Overall, P3b latency was reduced when participants actively engaged in a task-relevant context, suggesting faster processing under conditions of agency and relevance.

### 3.3. MVPA results

MVPA successfully dissociated left- and right-oriented gratings in the *Active Relevant*, *Passive Relevant*, and *Active Irrelevant* conditions. The largest significant temporal cluster was observed in the *Active Relevant* condition^1^, with a sustained significant difference going from 250 to 800 ms. A significant temporal cluster was also observed from 300 to 500 ms in the *Active Irrelevant* condition, and from 400 to 500 ms in the *Passive Relevant* condition. A repeated-measures ANOVA on maximum decoding scores revealed a significant main effect of agency (F(1,23) = 5.12, p = .033, η^2^*_G_* = 0.064), indicating that maximum classification accuracy was higher in the active compared to the passive conditions. By contrast, there was no significant main effect of relevance (F(1,23) = 0.05, p = .825) and no significant interaction (F(1,23) = 0.02, p = .881). However, this significant difference in maximum decoding scores between active and passive conditions did not remain when using alternative analysis parameters (e.g., 20 ms or 100 ms time windows, or different electrode selections).

### 3.4. EEG-behavior correlation results

#### 3.4.1. Faster reaction time from action-based prediction is correlated with shorter P3b latency

A significant positive relationship was found between the *Active Relevant* – *Passive Relevant* difference in RT and P3b latency (β = 450.63, SE = 182.36, t(22) = 2.47, p = .022, R² = .22, r = .47). Faster reaction times in the active condition relative to the passive condition (M = -20.58 ms; SD = -40.96) were associated with shorter P3b latencies for the same active – passive difference (M = -18.92 ms; SD = 42.37). In contrast, the relationship between P3b amplitude difference and RT difference was not significant (β = 6.27, SE = 3.67, t(22) = 1.71, p = .101, R² = .12, r = .34). This result suggests that faster behavioral responses under the *Active Relevant* condition were associated with corresponding changes in P3b latency, indicating a shared underlying process.

## 4. Discussion

A large number of studies have investigated how self-generated sensory events are processed and perceived. While some research has observed attenuation of self-generated outcomes (Dogge, Hofman, et al., 2019; Gentsch & Schütz-Bosbach, 2011; Roussel et al., 2013), a finding typically attributed to prediction mechanisms, other studies have found that stimuli generated by voluntary actions can be better perceived and even experienced with enhanced intensity (Desantis et al., 2014; Kiepe et al., 2023; Thomas et al., 2022). These contrasting findings raises the question of whether they reflect the same underlying predictive processes or are better explained by different mechanisms; specifically enhancement driven by attention and attenuation driven by prediction (Alink & Blank, 2021; Rungratsameetaweemana & Serences, 2019; Summerfield & Egner, 2016). Hence, the present study aimed to disentangle the contributions of attention and prediction to the processing of self-generated stimuli. To achieve this, we assigned each process to a distinct stimulus feature, i.e., color for attention and orientation for prediction. Specifically, participants viewed two consecutive gratings. The orientation of the first grating was either predicted by an action (*Active* condition) or not (*Passive* condition). Importantly, the color of the grating indicated whether a perceptual judgment was required (*Relevant* trials) or not (*Irrelevant* trials). Because relevant and irrelevant trials were equiprobable, the relevance cue, provided at the beginning of each block, contained no information about the likelihood of a feature’s stimulus (no specific color or orientation).

Our findings revealed that action-based predictions did not modulate early sensory processing, as indicated by the absence of N1a amplitude differences between active and passive conditions. However, later processing stages were modulated: the P3b amplitude was reduced for predicted/self-generated stimuli compared to unpredicted/externally-generated ones, but only when stimuli were task-relevant. Complementing the P3b results, MVPA analyses showed that despite this reduction in neural activity for predicted/self-generated events, the significant temporal clusters in which decoding accuracy was largest for predicted/self-generated and task-relevant stimuli. Additionally, P3b amplitudes were enhanced for relevant compared to irrelevant stimuli, but only when stimuli were unpredicted/externally-generated. Behavioral results further supported the P3b findings, as earlier P3b latencies were associated with shorter reaction times. Drift-diffusion modeling also indicated a tendency toward a higher drift rate and a smaller decision threshold for predicted/self-generated and task-relevant stimuli, compared to unpredicted/externally-generated but equally relevant ones.

### 4.1. Early sensory processing

Based on the crossover interaction between attention and prediction observed by Kok et al. (2012), we expected a larger N1a amplitude for predicted stimuli (*Active* condition) compared to unpredicted (*Passive* condition), when stimuli were relevant (i.e., required a perceptual response). Conversely, we expected the opposite effect for irrelevant stimuli, with a classical attenuation of the N1a for predicted stimuli. However, our results showed not impact of either prediction or attention on the N1a. These results diverge from studies (Marzecová et al., 2018), who reported an attentional gain effect on N1 component only for gratings with predicted orientation. They also contrast with findings from John-Saaltink et al. (2015), who observed brain activity reduction in primary sensory areas (measured by fMRI) for predicted compared to unpredicted gratings, but only when they were task-irrelevant. A key difference between those previous studies and the present experiment is twofold: here we manipulated feature-based attention rather than spatial attention, and predictions were generated through voluntary action rather than an external cue.

#### 4.1.1. Feature-based attention do not modulate early visual evoked potentials

Feature-based attention (FBA) has been shown to increase the firing rate of the neurons whose feature preference closely matches the attended feature value (e.g. the relevant color) (Gledhill et al., 2015, 2015; Liu et al., 2007; Martinez-Trujillo & Treue, 2004; McMains et al., 2007; Saenz et al., 2002). Nevertheless, little is known about the influence of FBA on ERPs. Its temporal onset varies across studies and recording sites, with the earliest effects occurring between 100–200 ms post-stimulus (Bondarenko et al., 2012; Eimer, 1997; Hayden & Gallant, 2005; He et al., 2004, 2008; Kenemans et al., 2002; W. Zhang & Luck, 2009) or between 200–300 ms (Gledhill et al., 2015; Moerel et al., 2022). Importantly, the FBA effect could depend on the prior knowledge about the specific relevant feature value (e.g., a particular color such as red). In the present study, because our aim was to disentangle predictive and attentional mechanisms, the relevance cue provided at the beginning of each block contained no information about the likelihood of a feature’s stimulus (it indicated the relevant color, but this color was equiprobable with the other one). Thus, participants could not predict the relevance of upcoming stimulus, thereby avoiding the preparatory aspect of top-down attention (Battistoni et al., 2017), which could be crucial for modulating early sensory processing (see Fannon & Mangun, 2013 for a review). Finally, in the current study, the absence of competing stimuli in the display could have further reduced the engagement of feature-based attention, even though stimulus competition is not indispensable (Bartsch et al., 2015).

#### 4.1.2. Visual action outcomes and action-based prediction do not modulate early visual evoked potentials

We did not observe a modulation of the active condition on the N1a amplitude. This result is in contrast with previous studies reporting sensory attenuation of the N1 component for predicted and/or self-generated stimuli compared to unpredicted and/or externally generated stimuli across various sensory modalities including vision (Baess et al., 2011; Bäß et al., 2008; Ghio et al., 2018; Klaffehn et al., 2019; Lange, 2009, 2011; Paraskevoudi & SanMiguel, 2023; Pinheiro et al., 2019; Poonian et al., 2015; Timm et al., 2013, 2014; Weiss et al., 2011; Vroomen & Stekelenburg, 2010; Gentsch et al., 2012; Gentsch & Schütz-Bosbach, 2011; Roussel et al., 2014; Hughes & Waszak, 2011).

This absence of an effect aligns with our MVPA results, which showed that information about grating orientation could only be decoded after the typical N1 time window (i.e., after ∼200 ms). Together, these findings suggest that, at least in our paradigm, the N1a (or even the P1; see Supplementary Material) may not index feature-specific processing such as orientation. Therefore, even if participants formed predictions about the grating’s orientation in the active condition, these predictions might not influence N1a activity because orientation information becomes more consistently represented only at later processing stages when prediction errors are used for model update (Bednark et al., 2015; Donchin, 1981; Polich, 2007), specifically during processes reflected by the P3b, where a modulation of the active condition was indeed found.

We can only speculate as to why orientation information was visible only during later processing stages. One possible explanation relates to the nature of the task. Participants were required to encode two stimulus features: color, which determined trial relevance, and orientation. It is plausible that participants prioritized color identification first to decide whether a response was needed, and only subsequently integrated orientation information. This task-dependent prioritization could explain why orientation-related activity emerged later in the processing stream.

Another possible explanation for the absence of N1 attenuation could be attributed to the relatively long delay between the action and its effect, which has been shown to reduce sensory attenuation in both the tactile and auditory domains (Bays et al., 2005; Horváth et al., 2012). However, we consider this explanation unlikely, since the delay between action and effect in our experiment remained constant and matched the interval used during the learning phase. Moreover, it appears that the temporal relationship between voluntary actions and their effects is learned quickly and remains stable once established (Dignath & Janczyk, 2017; Haering & Kiesel, 2012; Walsh & Haggard, 2013).

Another possible explanation is that the sensory attenuation observed in previous studies may not reflect prediction or self-generation per se, but rather a withdrawal of preparatory attention from predicted or self-generated events. Indeed, if we know in advance that what we are doing is irrelevant, we can disengage attentional resources from processing of our action outcome before its generation, resulting in reduced neural responses and potentially in an N1 attenuation. In our paradigm, such a mechanism could not occur, since the relevance of the upcoming stimulus was not known in advance. Participants had to attend to the visual outcome to determine its relevance based on its color, which could only be identified at stimulus onset. Therefore, they could not ignore the predicted orientation before stimulus onset, as they had no reason to do so – the predicted feature could prove either relevant or irrelevant depending on the stimulus color. As a result, preparatory attention likely remained equally engaged in both active and passive conditions, which may account for the absence of N1 attenuation in the active condition.

### 4.2. Later stages of processing

#### 4.2.1. Reduced decision evidence for self-generated outcomes when task-relevant

According to the synergistic interaction hypothesis mentioned in the introduction, we expected that if attention increases the gain of prediction error units (see Feldman & Friston, 2010), it should enhance model updating in higher-order processing areas, particularly when stimuli are task-relevant. Consistent with this idea, we observed a main effect of agency and significant interaction between attention and agency on the P3b component. Specifically, P3b amplitudes were overall reduced for self-generated compared to externally generated stimuli. Moreover, post-hoc analyses showed that the P3b was particularly reduced for predicted and relevant stimuli compared to unpredicted and equally relevant ones, whereas no difference was observed when stimuli were irrelevant.

Overall, our results align with the previous findings from the oddball literature, which show that the P3b component is sensitive to target probability and attentional resources (see Luck, 2014; Polich, 2011 for reviews). The P3b is thought to reflect the updating of high-level internal models or the storage of sensory representations in working memory in response to unpredicted or salient events (Donchin, 1981; Polich, 2007), a process that is modulated by task relevance when stimuli are linked to subsequent actions (Verleger et al., 2006). Therefore, our results could be explained by the notion that updating internal models is essential only for task-relevant features, those directly useful for task performance. This interpretation is also compatible with the idea that action-based predictions, when aligned with task goals, reduce the cognitive cost associated with evidence accumulation and the model updating (Zénon et al. 2019). In other words, participants may need to accumulate less evidence for stimulus categorization in the presence of relevant sensory predictions (see Darriba & Waszak, 2018; Kelly & O’Connell, 2013; O’Connell et al., 2012; Twomey et al., 2015 for the relationship between evidence accumulation and the P3b and Zénon et al., 2019 for work linking cognitive cost and predictive mechanisms). The MVPA results discussed below provide complementary information on the impact of predictions on the quality of sensory representation over time.

#### 4.2.2. Relevant action-based predictions optimize sensory representations and decision-making

In addition to ERP analyses, we performed Multivariate Pattern Analyses to decode the orientation of the first grating from EEG activity. Classification accuracy was significantly above chance level when the stimulus was relevant, predicted by action, or both. In contrast to previous studies (Kok, Jehee, et al., 2012; Yon et al., 2018), classification accuracy across time did not differ statistically between predicted and unpredicted stimuli or between relevant and irrelevant stimuli. The relatively low classification accuracy, likely influenced by the small size of the stimulus (7° of visual angle), its brief presentation duration (50 ms), and the presence of the colored circular noise patch, may have contributed to the absence of differences between conditions.

Interestingly, the reduction in P3b amplitude observed for predicted and relevant stimuli (compared to unpredicted and relevant ones), was accompanied by a prolonged window of significant decoding accuracy, the longest among all tested conditions. We speculated that action-based prediction and attention may not have enhanced the quality of sensory representation, but instead supported a more sustained and temporally extended neural representation of the stimulus, potentially facilitating down-stream decision-making processes.

This interpretation is consistent also with the behavioral results, which revealed no improvement in accuracy, but significantly faster response times for relevant and predicted/self-generated stimuli compared to relevant and unpredicted/externally-generated ones. The hierarchical Bayesian SSM revealed that this RT effect is possibly driven by the higher speed of evidence accumulation. This suggests that relevant action-based predictions may not enhance sensory representations (at least in our design), but rather optimize decision-making processes by accelerating perceptual decisions.

The P3b results may also reflect this optimization process. When stimuli are unpredicted/externally-generated, the need for evidence accumulation is greater, which can consequently slow down perceptual decisions. In contrast, relevant action-based predictions reduce uncertainty and thereby reduce the need for evidence accumulation. Since predicted events are already internally represented prior to stimulus presentation (Kok et al., 2017), decisions can be taken more efficiently, even if the quality of sensory evidence is comparable across active and passive conditions. In our case, this efficiency is reflected in faster decision times when predictions are available.

#### 4.2.3. Attentional gain effect only occurs for unpredicted/externally-generated stimuli

Based on previous reports of attentional gain effects (Lijffijt et al., 2009; Liu et al., 2007; Martinez-Trujillo & Treue, 2004; Saenz et al., 2002; Zinni et al., 2014), we expected larger P3b amplitudes for relevant compared to irrelevant stimuli. In line with this assumption, we observed a main effect of task relevance, with higher P3b amplitudes for relevant than irrelevant trials. However, post hoc comparisons showed that the relevance effect was driven by higher P3b amplitudes in the passive relevant condition compared to passive irrelevant condition, whereas no difference was observed between the active conditions. This suggest that observers accumulated more sensory evidence when stimuli where relevant to the ongoing task, particularly in the passive condition where predictive information from self-generated actions was absent.

Overall, the P3b results align with previous studies (Smout et al., 2019, 2020), supporting the notion, within predictive coding theory, that attention optimizes the precision (i.e. reliability) of ascending prediction error (PE) signals by boosting synaptic gain of PE units (Feldman & Friston, 2010; Kanai et al., 2015). In this framework, if attention enhances PE signals, it implies that the more unexpected a stimulus is, the larger the number of produced PEs, leading to stronger model updating in higher-order processing areas that give rise to the P3b response. Accordingly, a higher P3b amplitude observed in the passive relevant trials compared to active relevant and passive irrelevant trials may reflect enhanced updating when predictions are absent and stimuli are task-relevant (i.e., attended).

## 5. Conclusion

In conclusion, we have found that action-based prediction and top-down attention both modulate late processing stages and interact with one another. Specifically, when aligned with behavioral goals, action-based predictions appear to reduce the amount of evidence needed and increase the rate of its accumulation, while maintaining a similar sensory representation quality as in the absence of prediction and action. This pattern suggests an optimization process that enhances processing efficacy without compromising internal representation.

Our findings on late perceptual processing support extended predictive coding models in which attention globally boosts the synaptic gain of prediction error units. However, we found no modulation of early sensory processing – regardless of their behavioral relevance. This null effect challenges motor control and predictive coding theories which propose early sensory attenuation of predicted or self-generated sensory inputs – either through cancellation or sharpening mechanism. Further studies aiming at isolating the influence of predictive and attentional mechanisms are needed to clarify this issue. In particular, early sensory effects may depend on the presence of preparatory attention. Such efforts, alongside investigations across other sensory modalities and comparisons between action-based and externally cued predictions may provide a more precise understanding of how attention, prediction and voluntary action jointly contribute to sensory and perceptual processing.

## CRediT authorship contribution statement

**Thomas Holstein:** Conceptualization, Data curation, Formal analysis, Investigation, Methodology, Project administration, Software, Visualization, Writing – original draft, Writing – review & editing. **Jean-Christophe Sarrazin**: Conceptualization, Data curation, Methodology, Resources, Supervision, Writing – review & editing. **Bruno Berberian:** Conceptualization, Data curation, Methodology, Supervision, Writing – review & editing. **Andrea Desantis:** Conceptualization, Data curation, Funding acquisition, Methodology, Supervision, Writing – review & editing.

## Appendix. Supplementary Material

## Acknowledgements

We are grateful to Patrick Haggard for his comment and suggestions on the design of the task.

This research was funded by the Agence Nationale de la Recherche, research funding ANR JCJC, project number ANR-18-CE10-0001, awarded to Andrea Desantis.

## Footnotes

1 Note that these results are stable across different analysis parameters. For example, the significant temporal cluster was larger for the active conditions than the passive conditions across other time windows tested (20 ms and 100 ms time windows) and when selecting different numbers of electrodes (pools of 5 or 15 electrodes per time window, posterior electrodes, or all electrodes). See Supplementary Material.

