## Appendix - Supplementary Material for "Interplay of action-based prediction and top-down attention: EEG evidence for joint modulation of late perceptual processing"

#### 1. Behavioural analyses

Since only the *Relevant* conditions required a response, behavioral analyses were performed only for these conditions. Specifically, we calculated sensitivity ($\boldsymbol{d'}$) and response bias ($\boldsymbol{c}$) under *Active Relevant* and *Passive Relevant* conditions (Green & Swets, 1966; Hautus et al., 2021). A clockwise rotation of the second stimulus was defined as the presence of the signal, while an anticlockwise rotation indicated its absence.

Sensitivity ($\boldsymbol{d'}$) quantifies the separation between signal (i.e. clockwise rotation) and noise (i.e. anticlockwise rotation), reflecting participants' ability to discriminate stimulus orientation, independent of response bias (Juravle & Spence, 2011). Response bias ($\boldsymbol{c}$) measures the tendency of the participants to favor one response over another when judging the orientation of the second grating. A positive criterion $\boldsymbol{c}$ indicates a conservative bias, requiring stronger evidence to respond "clockwise", leading to fewer false alarms but more misses. Conversely, a negative $\boldsymbol{c}$ indicates a liberal bias, requiring less evidence to respond "clockwise," leading to more hits but also more false alarms. Sensitivity and response bias were calculated as follows:

$$d' = Z(Hit\ rate) - Z(False\ alarm\ rate)$$
$$c = -\frac{1}{2}[Z(Hit\ rate) + Z(False\ alarm\ rate)]$$

The paired t-tests on $\boldsymbol{d'}$ and $\boldsymbol{c}$ indicated no significant difference ($\boldsymbol{d'}$ : p = .280 ; $\boldsymbol{c}$ : p = .570).

#### 2. Hierarchical Drift Diffusion Model (HDDM)

The HDDM sampling was performed with 2000 draws per chain (8000 effective samples total) and 2000 warm-up tuning iterations across four chains (target acceptance = 0.95; max tree depth = 12) and with a non-centered parametrization, ensuring convergence and robust posterior estimation. Convergence was assessed using rank-normalized $\hat{R}$ and effective sample size (ESS) diagnostics. The maximum $\hat{R}$ across all monitored parameters was 1.009, below the recommended threshold of 1.01, indicating adequate convergence. The minimum bulk-ESS (287) and tail-ESS (714) exceeded conventional reliability criteria (bulk-ESS > 100, tail-ESS > 100), confirming aceptable sampling efficiency across parameters. No divergent transitions were observed, and trace plots indicated good mixing across chains. These results provide strong evidence that the model converged satisfactorily and that posterior inferences are robust.

### 3. EEG-behavior correlation analysis

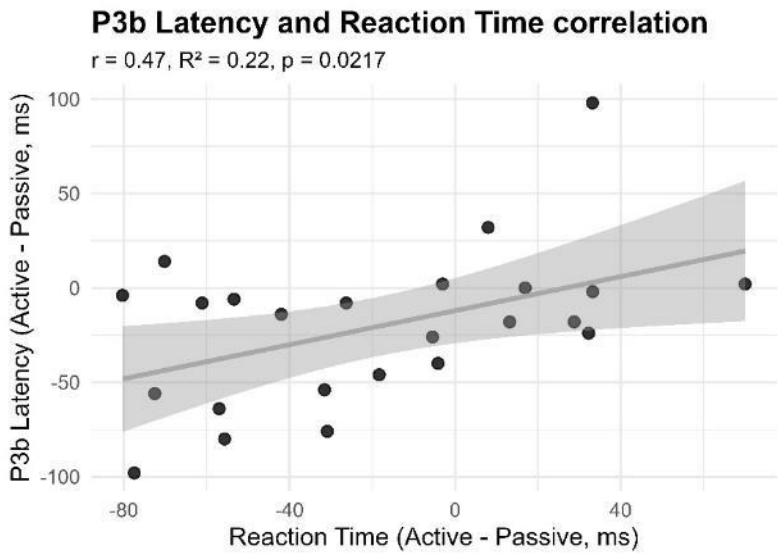


***Figure 1:*** *Scatter plot illustrating the significant linear correlation ($r = 0.47$; $R^2 = 0.22$; $p = .022$) between the change in P3b latency (Active – Passive, in µV) and the corresponding difference in reaction times (Active – Passive, in milliseconds) across participants. Each point represents an individual participant, while the regression line (±95% confidence interval, shaded area) depicts the best-fitting linear model.*

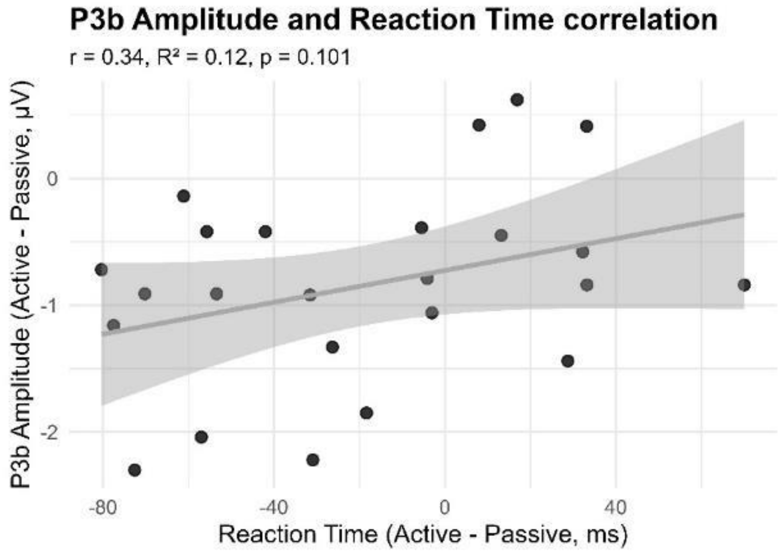


***Figure 2:*** *Scatter plot illustrating the non-significant linear correlation ($r = 0.34$; $R^2 = 0.12$; $p = .101$) between the change in P3b amplitude (Active – Passive, in µV) and the corresponding difference in reaction times (Active – Passive, in milliseconds) across participants. Each point represents an individual participant, while the regression line (±95% confidence interval, shaded area) depicts the best-fitting linear model.*

### 4. EEG preprocessing

All preprocessing parameters are detailed in Table 1. On average, 3.44% of epochs were rejected per participant (SD = 3.37%).

### 5. ERP analysis

#### 1.1. P1 analysis

The time-window for our analysis of this component was set at 80–200 ms. Some studies indicate that this component is associated with predictive or attentional mechanisms, showing enhanced P1 for spatially or temporally predicted stimuli (Doherty et al., 2005; Hughes & Waszak, 2011). We planned to conduct statistical analyses on Oz and POz electrodes, but visual inspection on ERP grand average did not reveal any P1 waveforms patterns. Consequently, we conducted statistical analyses only on the electrode where the maximum positive deflection in the time window of interest was reached in the grand average – in P6 electrode. Indeed, P1 component is often largest at lateral occipital electrode sites, with a peak latency of approximately 100 ms (Luck, 2014).

#### 1.2. Null effect on P1

For P1 component on P6 electrode – where the maximum positive deflection in the time window of interest (80-200ms) was reached – a repeated measure ANOVA with agency (active, passive) and task-relevance (relevant, irrelevant) as factors revealed no significant main effect of agency ($F(1,23)=0.04$, $p=.834$) and task-relevance ($F(1,23)=1.22$, $p=.280$) and no significant interaction ($F(1,23)=0.60$, $p=.445$) (Supp. Mat. Fig. 1).

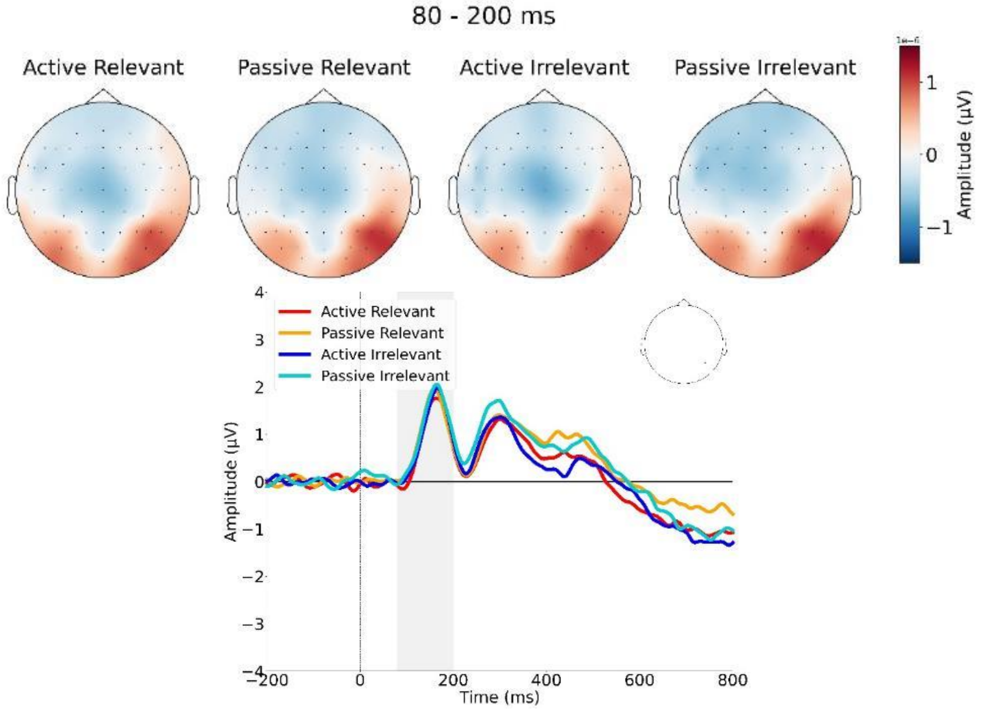

***Figure 1:*** *(top) Topography of brain activity averaged over the P1 component time window (80–200 ms). (bottom) Grand average ERPs of the P1 component on P6 electrode.*

##### 1.3. N170 analysis

The latest N1 subcomponent – the N170 – peaks 150-200 ms post-stimulus at posterior electrode sites. N170 could be an index of mismatches between prior expectations and actual stimulus input in higher-level vision (Baker et al., 2021; Johnston et al., 2017; Robinson et al., 2020) with an enhanced N170 for unpredicted stimuli. However, visual inspection of ERP grand average on time-window 150-200ms at central and lateral posterior occipital electrodes and parietal electrodes did not reveal any N170 waveforms patterns, so statistical analyses for this component were discarded. This may be due to the absence of prediction violations in this experimental paradigm. Indeed, there were only fully 100% congruent predictions (active condition) or an absence of prediction (passive condition) with equiprobability of a stimulus oriented to the left or right direction.

##### 1.4. P2 analysis

The P2 component is the second major positive peak overall and is observed at approximatively 200 ms. The functional significance of the P2 remains relatively poorly understood. Growing evidence suggests that the P2 may reflect stimulus encoding mechanisms, including processes of selective attention, feature detection, and high-order perceptual processing (Dunn et al., 1998; Evans & Federmeier, 2007; Geisler & Murphy, 2000; Luck & Hillyard, 1994; Sugimoto & Katayama, 2013). Interestingly, the P2 has been linked to cognitive control and inhibitory processing (Barry & De Blasio, 2013; Freunberger et al., 2007; Ghin et al., 2022; Johnstone et al., 2005; Kühn et al., 2009; Senderecka et al., 2012; Stock et al., 2016). In parallel, other research has explored the link between P2 and the sense of agency, with mixed findings. Some studies report increased P2 amplitudes when participants perceive greater agency over auditory stimuli (Timm et al., 2016), whereas others find smaller P2 amplitudes when participants control the timing of stimuli (Harrison et al., 2021).

Consequently, we conducted statistical analyses in all midline electrodes where the P2 waveform pattern was present in the grand average. This led us to focus on Pz, POz and Oz electrodes in the time-window 190-295 ms and it was on POz electrode that the maximum positive deflection was reached.

##### 1.5. P2 is modulated by task-relevance but not action-based prediction

In the time window of P2 component (190–295 ms post-stimulus onset), the repeated measure ANOVA revealed a significant main effect of task-relevance on Oz electrode ($F(1,23) = 14.24$, $p < .001$, $\eta_G^2 = 0.005$) and POz electrode ($F(1,23) = 11.49$, $p = .003$, $\eta_G^2 = .005$) which was the electrode where the maximum positive deflection was reached for this component (Supp. Mat. Fig. 2). Irrelevant stimuli elicited a larger P2 peak (M = 6.93 µV, SD = 3.54 on Oz electrode) than relevant stimuli (M = 6.44 µV, SD = 3.63 on Oz electrode). No significant main effect of agency was observed ($F(1,23)=1.31$, $p=.265$, for Oz electrode; $F(1,23)=1.76$, $p=.198$, for POz electrode). The interaction between task-relevance and action-based prediction was not significant on both electrodes (F = 1.94, p = .177, and F = 1.44, p = .242, for respectively Oz and POz electrodes).

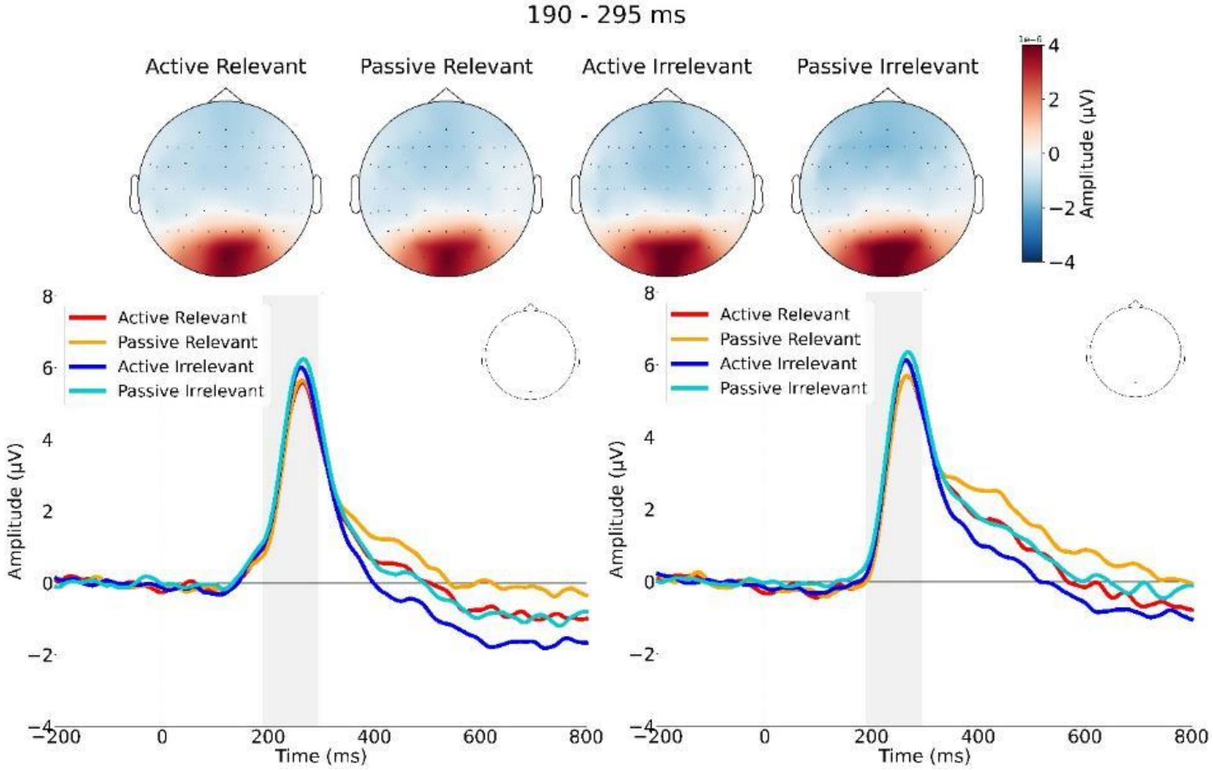


***Figure 2:*** *(top) Topography of brain activity averaged over the P2 component time window (190–295 ms). (bottom) Grand average ERPs of the P2 component on Oz (left) and POz (right) electrodes.*

#### 1.6. N2 analysis

The N2 component is the second major negative peak and is observed between 200-350 ms. It can be described in three subcomponents: N2a, N2b, N2c. The N2a is an automatic effect that occurs for auditory mismatches, even if they are task-irrelevant. This effect is more commonly known as the mismatch negativity (MMN) but it is mostly specific to auditory stimuli (see Garrido et al., 2009 for a review). N2b is the anterior sites component so it is also called anterior N2, and N2c is the posterior sites component so it is also called posterior N2. The N2b is related to cognitive control notably response inhibition typically observed in the go/no-go paradigm (see Folstein & Van Petten, 2008 for a review). When the no-go stimulus is presented (i.e. when the participant is not required to response to the stimulus), N2b is largest than if it is the go stimulus (i.e. when the participant is required to response). This enhanced effect is particularly pronounced when the go stimulus is more common than the no-go stimulus. As our paradigm includes an equiprobable go/no-go, we conducted analyses on this component in all midline anterior electrodes from AFz to Cz where the maximum negative deflection was reached, and thus also including FCz and Fz. The time-window was set at 200-350 ms.

The functional significance of the N2c component is less clear than for N2b. It has been shown that N2c is highly sensitive to the probability of the target (Luck & Hillyard, 1994). According to some authors, it may reflect the degree of attention required for processing stimuli in visual cortex (Suwazono et al., 2000). Another similar component occurring at approximately the same time interval is the N2pc (N2-posterior-contralateral) component, which is observed at posterior scalp sites contralateral to an attended object and reflects some aspect of the focusing of attention. As there is no lateralized stimulus in our current paradigm, we did not conduct analyses on the N2pc component. Moreover, visual inspection of ERP grand average on time-window 200-350 ms at all midline posterior electrodes did not reveal any N2c waveforms patterns, so statistical analyses for this component were discarded.

##### 1.7. N2b show motor inhibition when motor system was activated by the previous action-based prediction

For N2b component (200–350 ms post-stimulus onset) on Cz and FCz electrodes, we applied a Yeo-Johnson transformation to all conditions (Yeo & Johnson, 2000) as the data did not follow a normal distribution (Shapiro-Wilk test, $p < .05$). On Cz electrode, the two-way rANOVA revealed a significant main effect of action-based prediction ($F(1,23)=8.39$, $p=.008$, $\eta_G^2=.024$), no effect of task-relevance ($F(1,23)=.92$, $p=.346$) and a significant interaction ($F(1,23)=10.20$, $p=.004$, $\eta_G^2=.013$). The N2 was larger in active conditions (M = -2.94 µV, SD = 1.43) than in passive conditions (M = -2.70 µV, SD = 1.59). Subsequent post-hoc paired t-tests with Bonferroni correction identified significant difference between active irrelevant condition and passive irrelevant condition ($t(23) = -3.80$, $p =.0055$, $d = -0.776$) (Supp. Mat. Fig. 3).

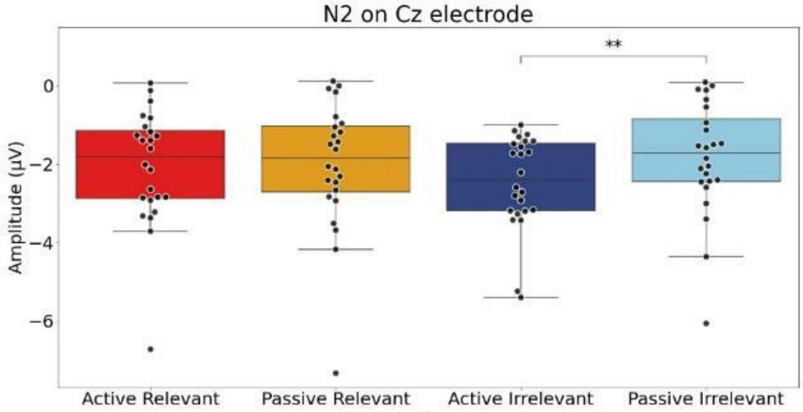


***Figure 3:*** *Boxplots of N2b peaks amplitude on Cz electrode illustrating the post-hoc comparisons with a significant difference between the active irrelevant condition and the passive irrelevant condition.*

On FCz electrode, the repeated measure ANOVA revealed no significant main effect of action-based prediction ($F(1,23)=3.49$, $p=.074$), no significant main effect of task-relevance ($F(1,23)=.89$, $p=.355$), but a significant interaction ($F(1,23)=6.61$, $p=.017$, $\eta_G^2=.007$) (Supp. Mat. Fig. 4). Subsequent post-hoc paired t-tests with Bonferroni correction showed that the difference between active irrelevant condition and passive irrelevant condition approached statistical significance ($t(23) = -2.83$, $p =.0573$, $d = -0.577$).

On Fz and AFz electrode the two-way rANOVA revealed no significant main effect of action-based prediction ($F(1,23)=0.05$, $p=.832$ and $F(1,23)=0.03$, $p=.865$ respectively) and task-relevance ($F(1,23)=1.10$, $p=.305$ and $F(1,23)=0.11$, $p=.745$ respectively) and no significant interaction ($F(1,23)=0.00$, $p=.986$ and $F(1,23)=0.55$, $p=.465$ respectively) (Supp. Mat. Fig. 4).

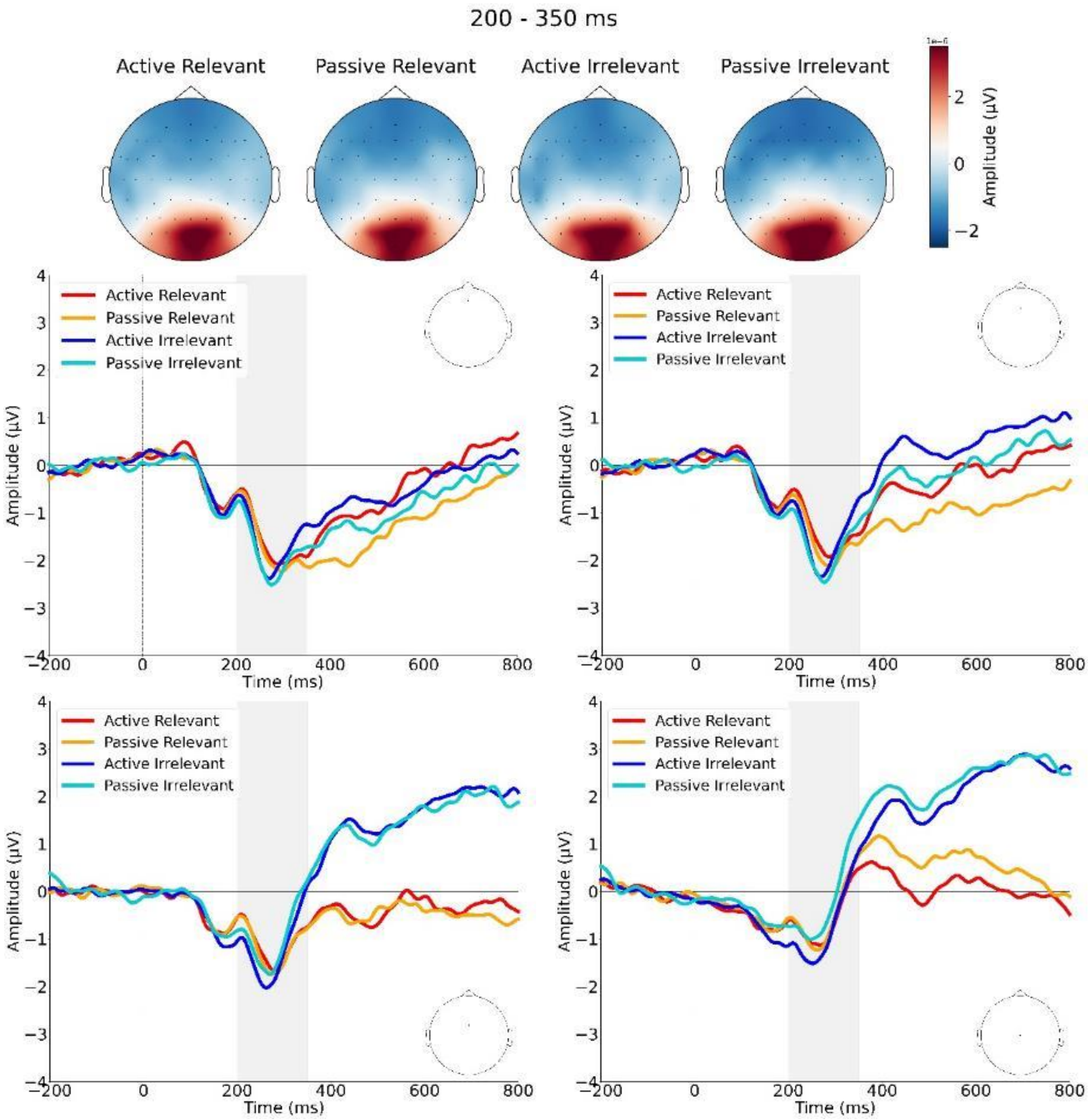


***Figure 4:*** *(top) Topography of brain activity averaged over the N2b component time window (200–350 ms). (bottom) Grand average ERPs of the N2b component on AFz (upper left), Fz (upper right), FCz (lower left) and Cz (lower right) electrodes.*

#### 1.8. P3 analysis

For Cz and CPz electrodes we used the time-windows 300-500ms because the maximum peak was sometimes reached just before 350ms, and because there was no P2 component peak to be confused with the P3 peak for these two electrodes.

#### 1.9. P3 is modulated by task-relevance and action-based prediction

The two-way rANOVA for the P3 component on Cz electrode revealed a significant main effect of action-based prediction (F(1,23) = 6.89, p = .015, $\eta_G^2$= .025) and task-relevance (F(1,23) = 13.95, p = 0.001, $\eta_G^2$= .119), but no significant interaction (F(1,23)=0.73, p=.400) (Supp. Mat. Fig. 5). On this electrode, the direction of the effect of task relevance was opposite to the direction of the effect on Pz electrode. Here, P3 peak amplitude was larger for irrelevant conditions (M = 3.95 µV, SD = 1.72) than for relevant conditions (M = 3.75 µV, SD = 1.40). The P3 peak amplitude was larger for passive conditions (M = 2.41 µV, SD = 1.44) than for active conditions (M = 2.02 µV, SD = 1.76). On CPz electrode, the two-way rANOVA revealed a significant main effect of action-based prediction (F(1,23)=12.10, p=.002, $\eta_G^2$=.083) but no significant main effect of task-relevance (F(1,23)=.89, p=.355) and no significant interaction (F(1,23)=3.31, p=.082) (Supp. Mat. Fig. 5).

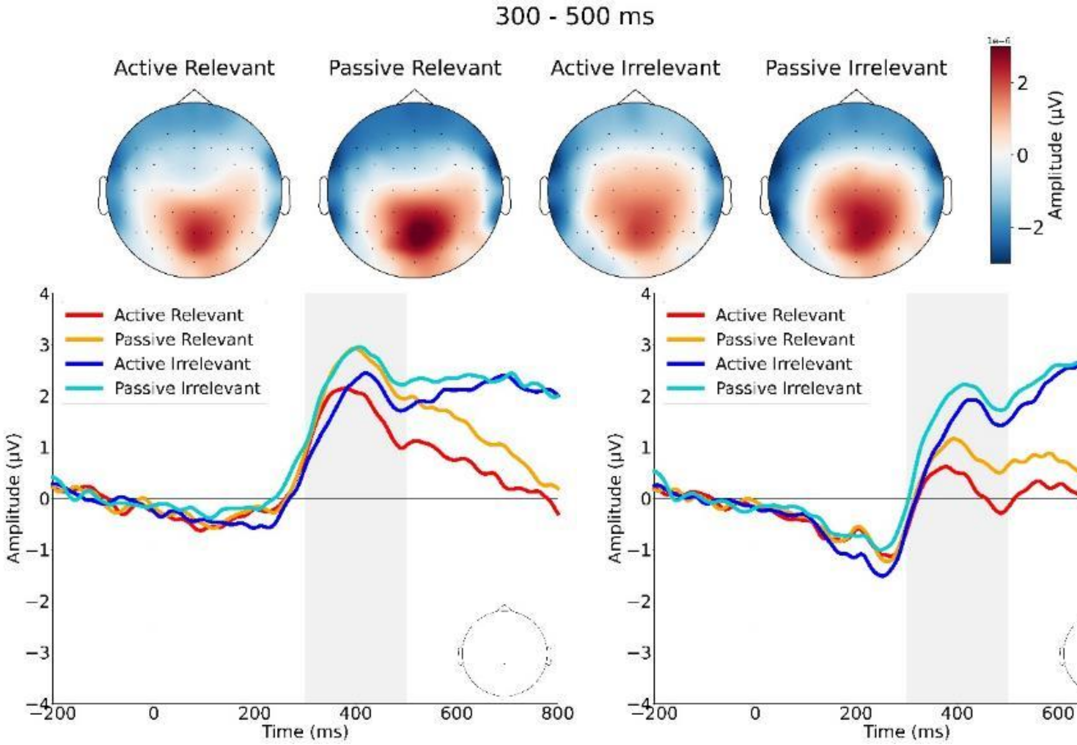


***Figure 5:*** *(top) Topography of brain activity averaged over the P3 component time window (350–500 ms). (bottom) Grand average ERPs of the P3 component on CPz (left) and Cz (right) electrodes.*

### 6. MVPA analysis

MVPA analysis with other time windows and electrode selections than those presented in the main article further support the result showing that the significant temporal cluster was consistently largest in the active relevant condition compared to any other condition. Indeed, results are relatively stable across different analysis parameters such as alternative time windows (20, 50 or 100 ms), and different electrode selection strategies (pools of 5 or 15 electrodes per time window, posterior electrodes, or all electrodes) (Supp. Mat. Fig. 6-19). When classification accuracy was significantly above chance, no condition showed superior performance compared to others.

To ensure that the EEG signals of the motor action itself did not induce prolonged changes that could bias the MVPA classification, we examined event-related potentials with a pre-action baseline (from -200 to 100 ms before action). We confirmed that after the motor response the EEG signal stabilized, indicating that the classification should not be driven by action-related activity associated with the orientation of grating 1 (Fig. 20).

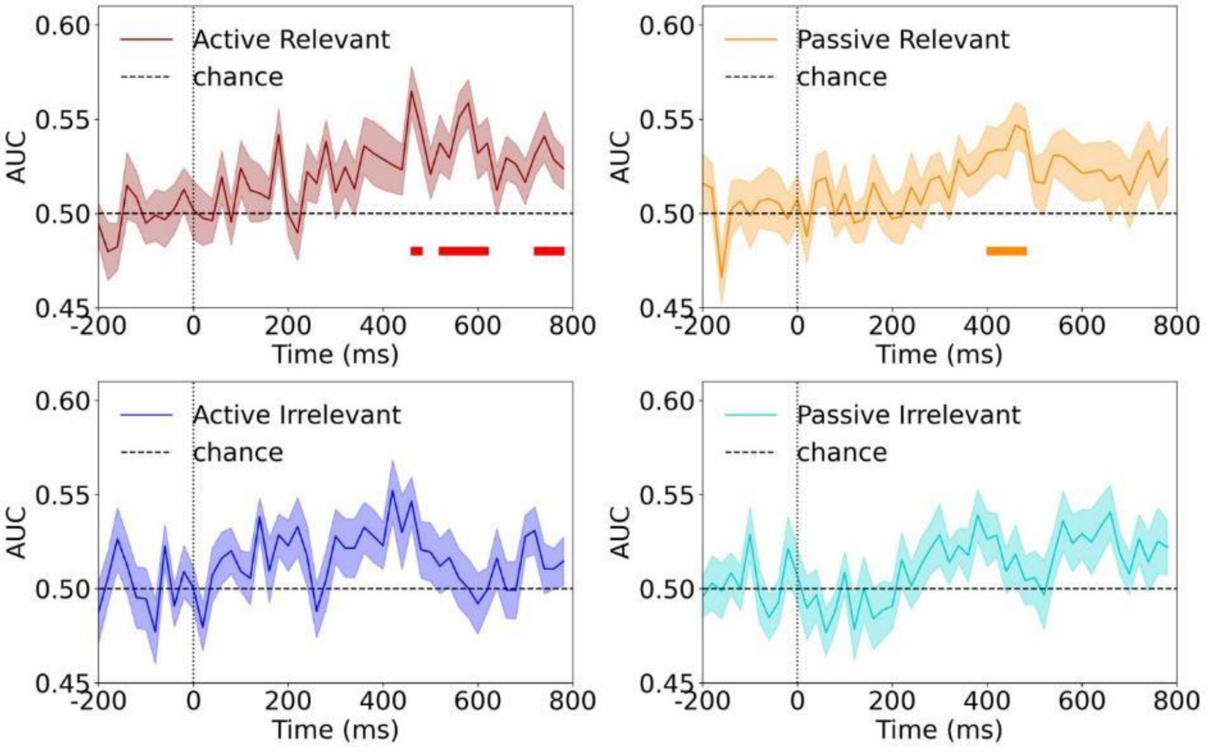


***Figure 6:*** *AUC scores over time for the 20 ms time window classifier with 5 electrodes per time point selection.*

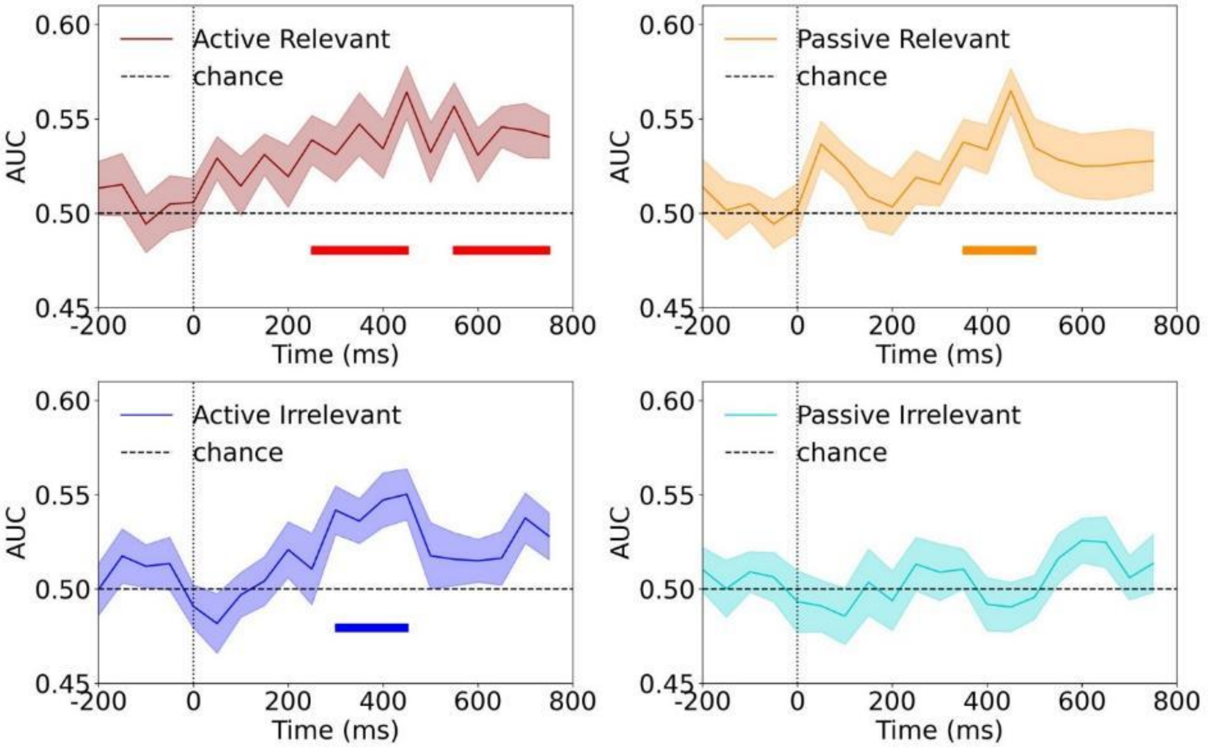

***Figure 7:*** *AUC scores over time for the 50 ms time window classifier with 5 electrodes per time point selection.*

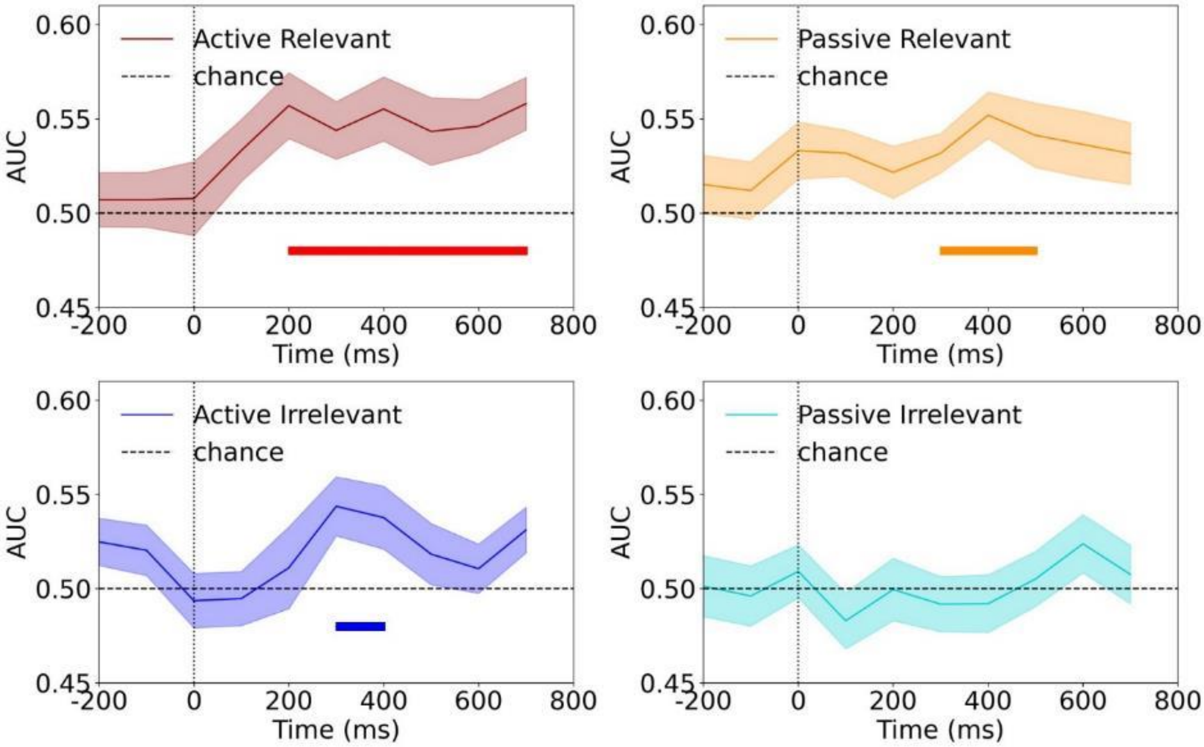


***Figure 8:*** *AUC scores over time for the 100 ms time window classifier with 5 electrodes per time point selection.*

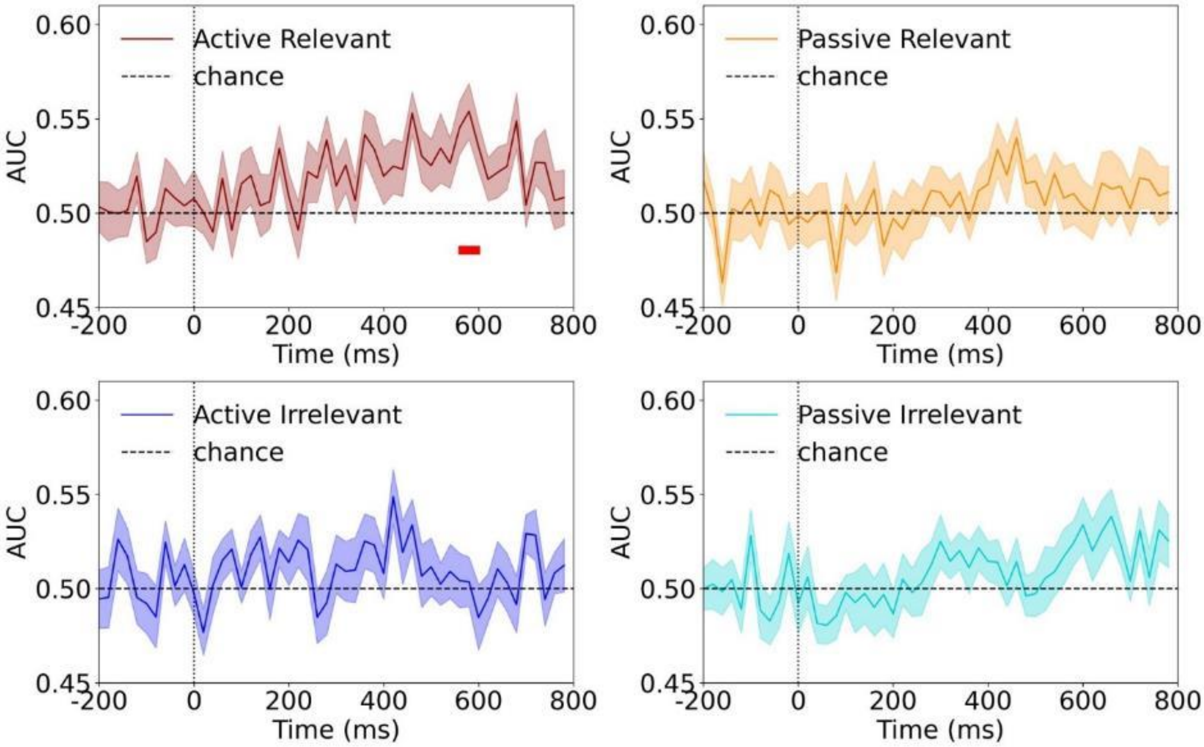


***Figure 9:*** *AUC scores over time for the 20 ms time window classifier with 10 electrodes per time point selection.*

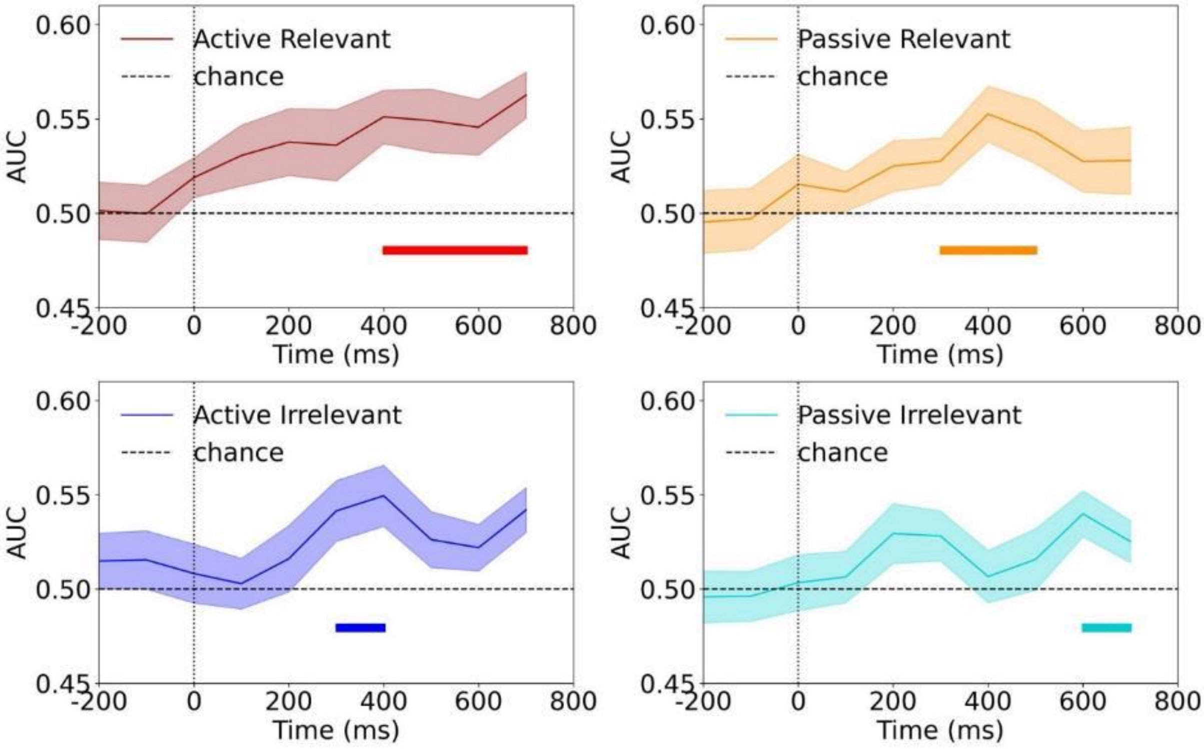


***Figure 10****: AUC scores over time for the 100 ms time window classifier with 10 electrodes per time point selection.*

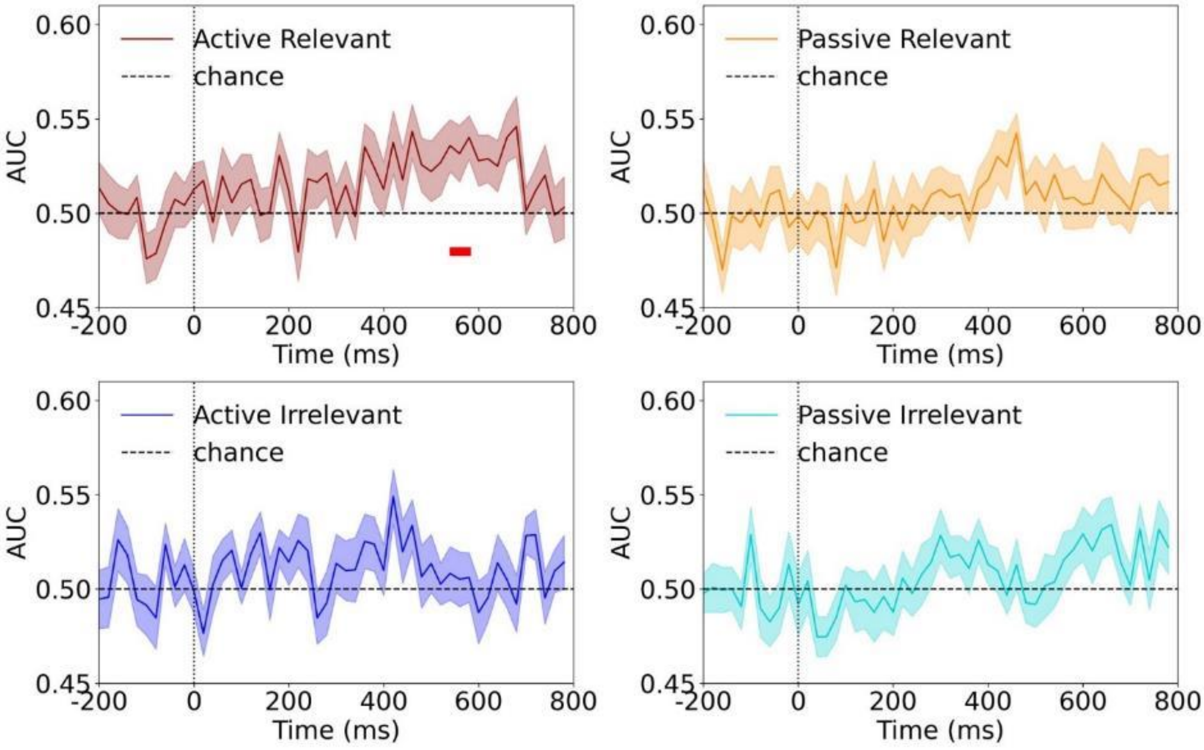


***Figure 11:*** *AUC scores over time for the 20 ms time window classifier with 15 electrodes per time point selection.*

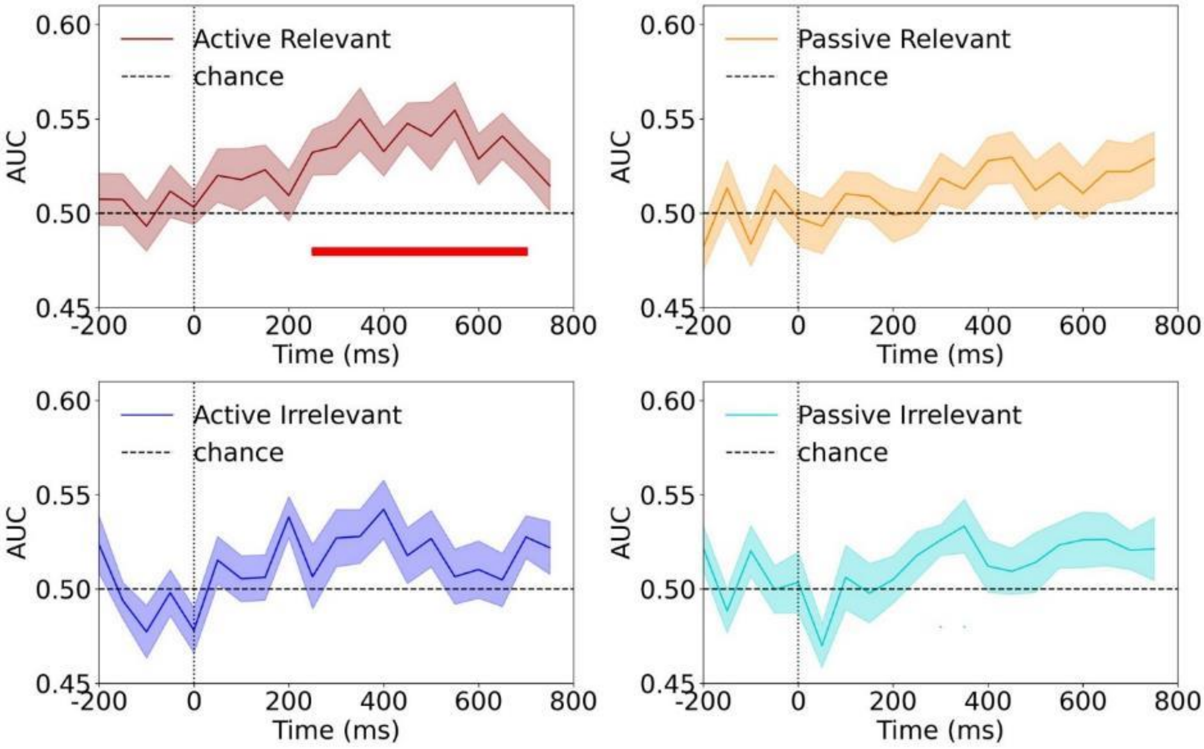


***Figure 12:*** *AUC scores over time for the 50 ms time window classifier with 15 electrodes per time point selection.*

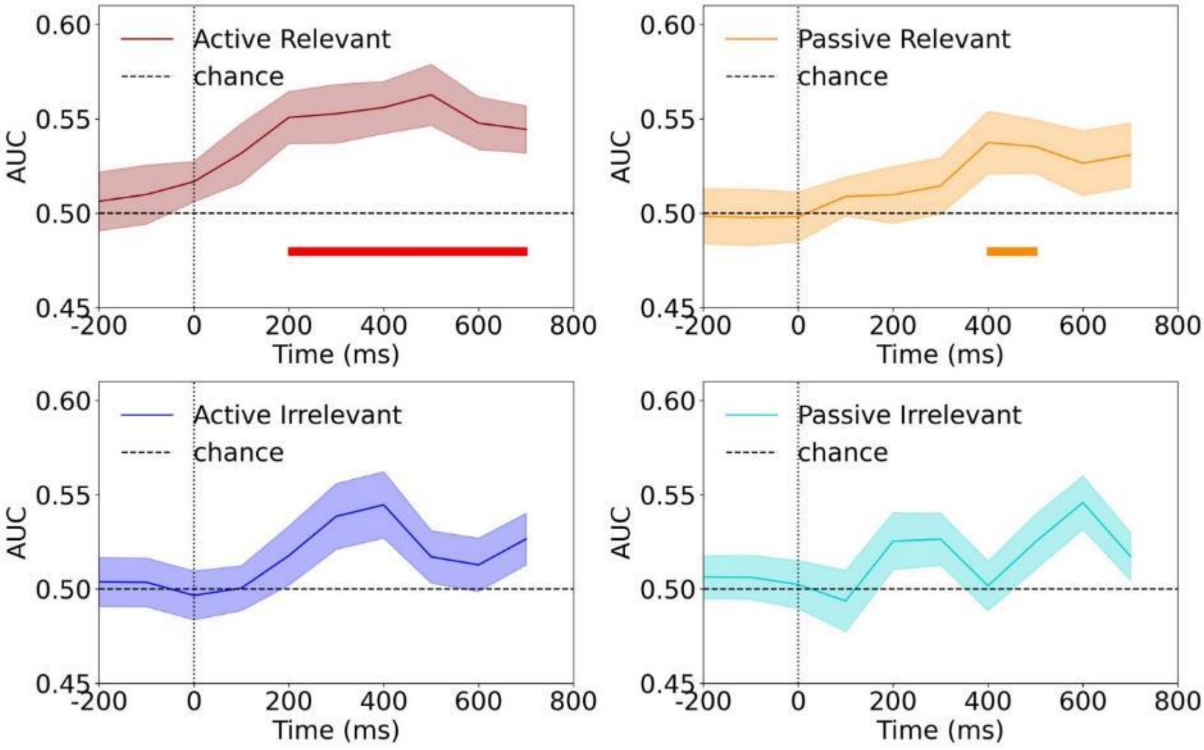


***Figure 13:*** *AUC scores over time for the 100 ms time window classifier with 15 electrodes per time point selection.*

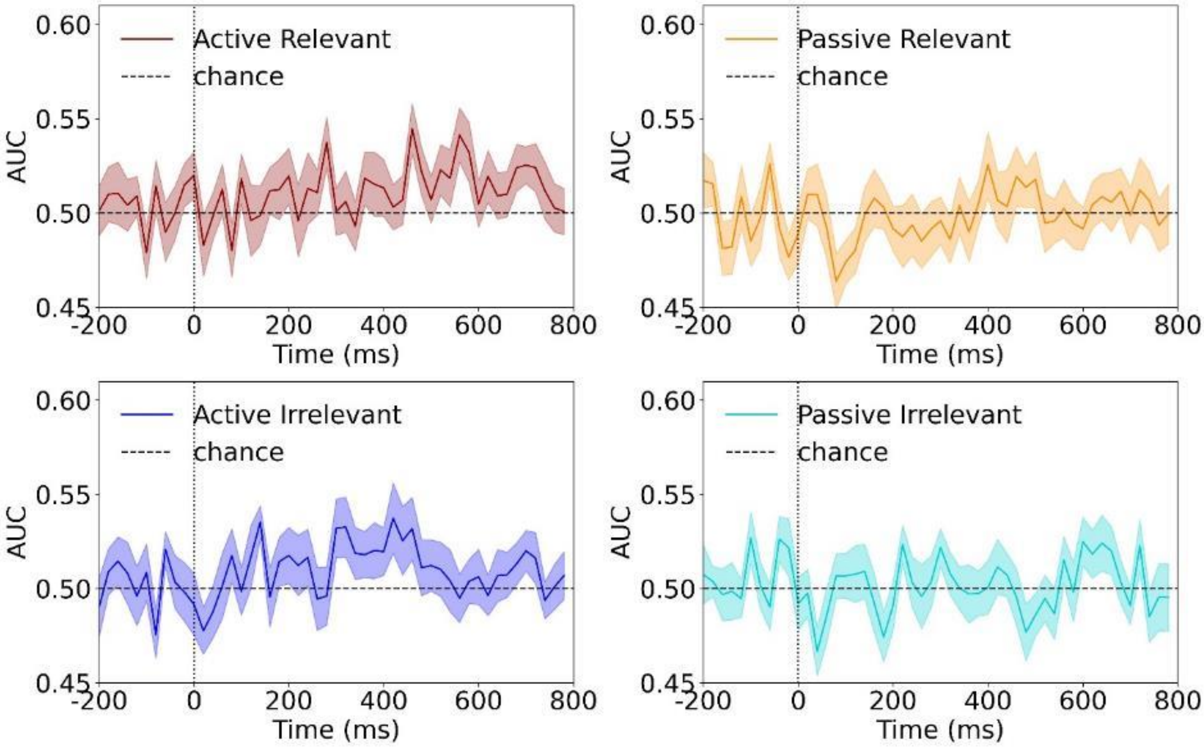


***Figure 14:*** *AUC scores over time for the 20 ms time window classifier with posterior electrodes selection (Oz, O1, O2, Pz, P1, P2, P3, P4, P5, P6, P7, P8, POz, P03, PO4, PO7, PO8, TP7, CP5, CP3, CP1, CPz, CP2, CP4, CP6, TP8, TP9, TP10).*

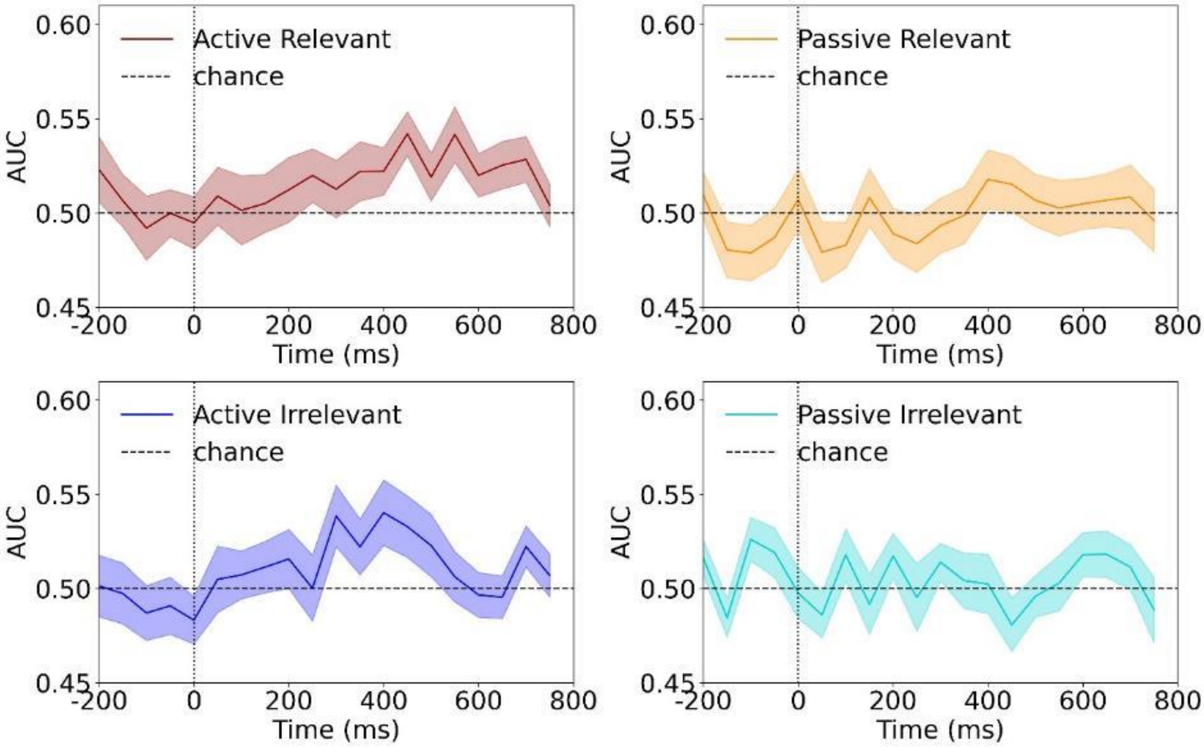


***Figure 15:*** *AUC scores over time for the 50 ms time window classifier with posterior electrodes selection (Oz, O1, O2, Pz, P1, P2, P3, P4, P5, P6, P7, P8, POz, P03, PO4, PO7, PO8, TP7, CP5, CP3, CP1, CPz, CP2, CP4, CP6, TP8, TP9, TP10).*

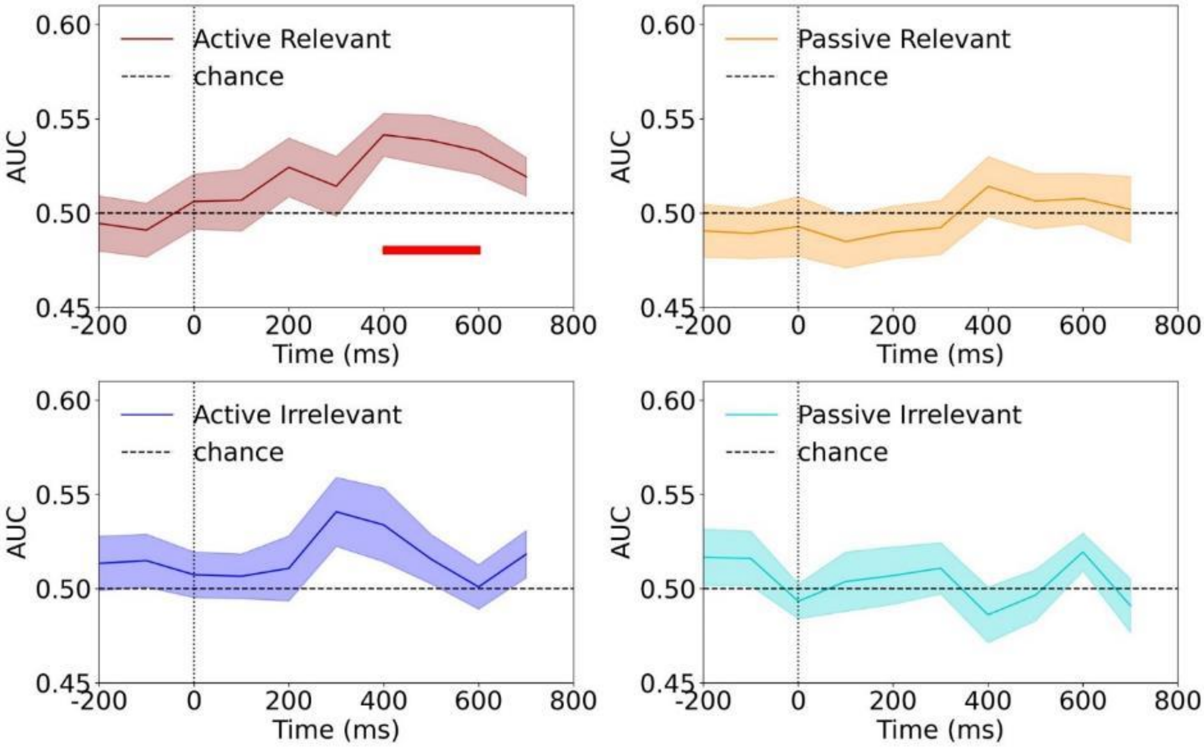


***Figure 16:*** *AUC scores over time for the 100 ms time window classifier with posterior electrodes selection (Oz, O1, O2, Pz, P1, P2, P3, P4, P5, P6, P7, P8, POz, P03, PO4, PO7, PO8, TP7, CP5, CP3, CP1, CPz, CP2, CP4, CP6, TP8, TP9, TP10).*

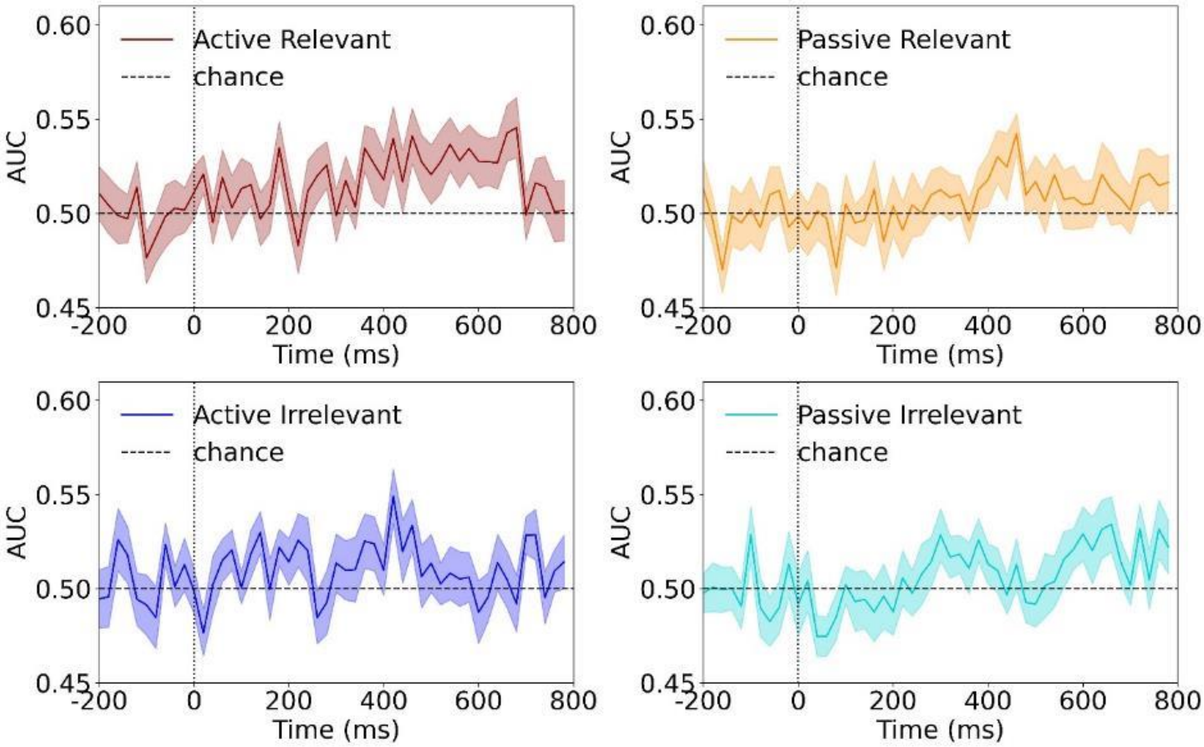


***Figure 17:*** *AUC scores over time for the 20 ms time window classifier with all electrodes selected.*

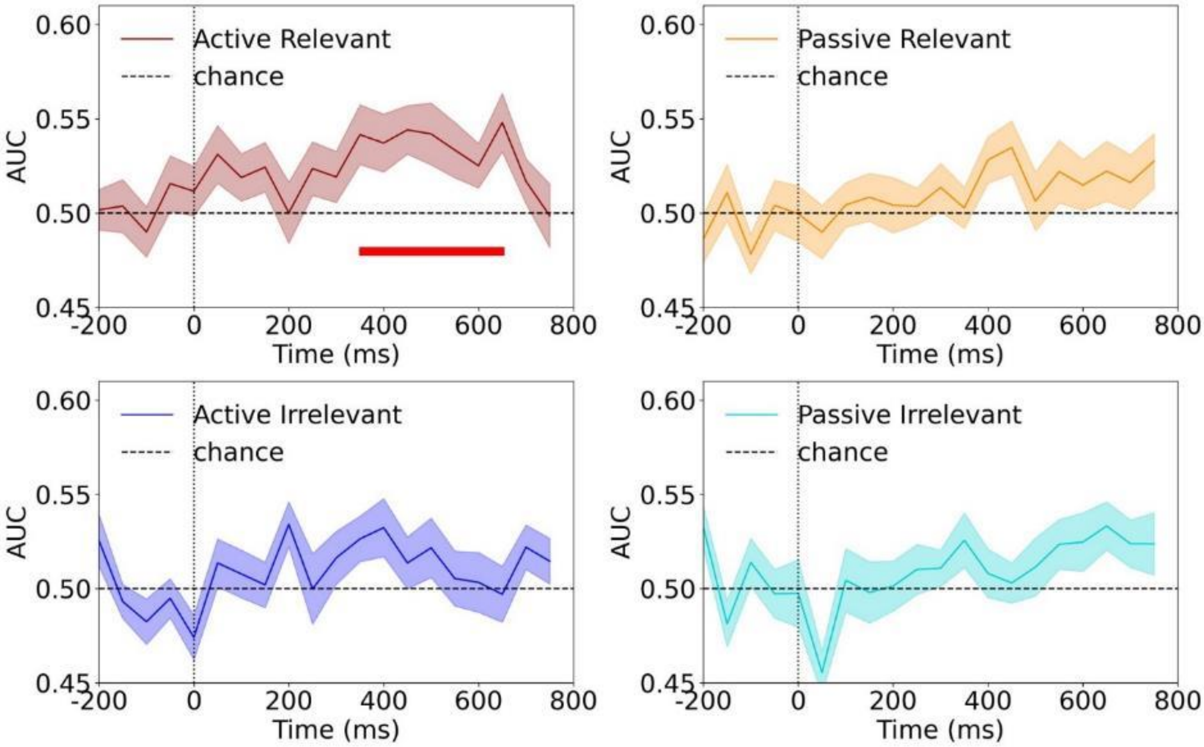


***Figure 18:*** *AUC scores over time for the 50 ms time window classifier with all electrodes selected.*

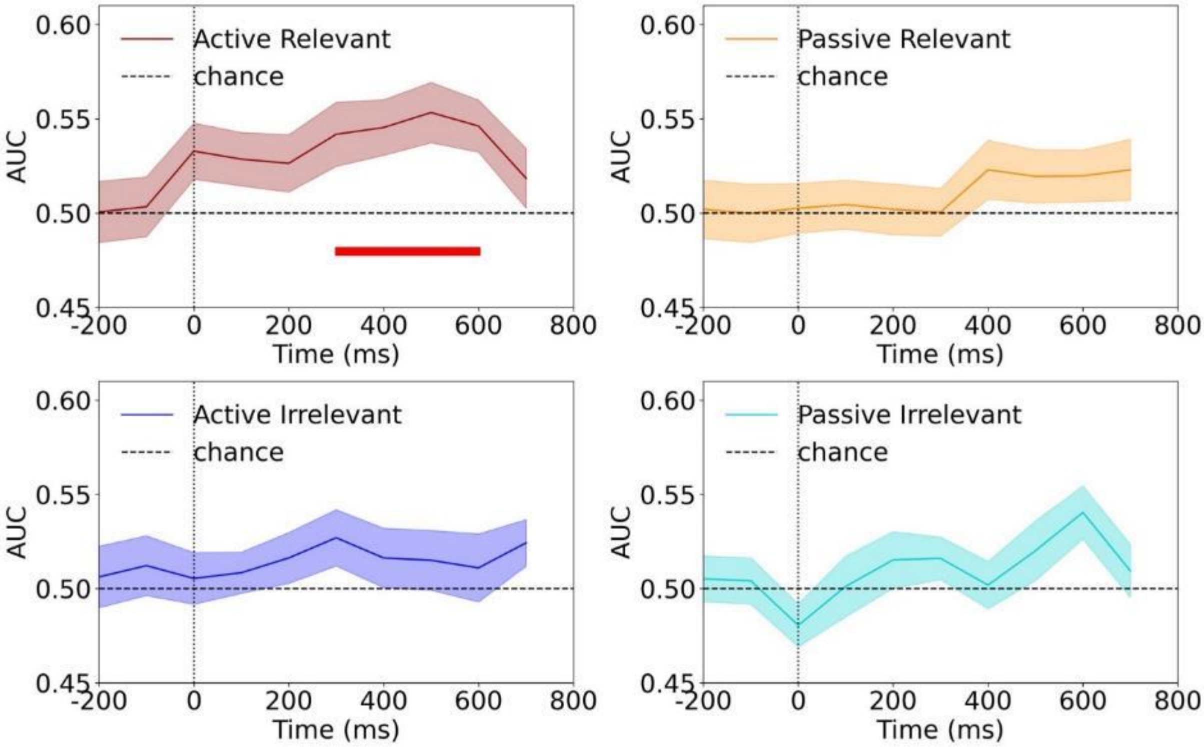


***Figure 19:*** *AUC scores over time for the 100 ms time window classifier with all electrodes selected.*

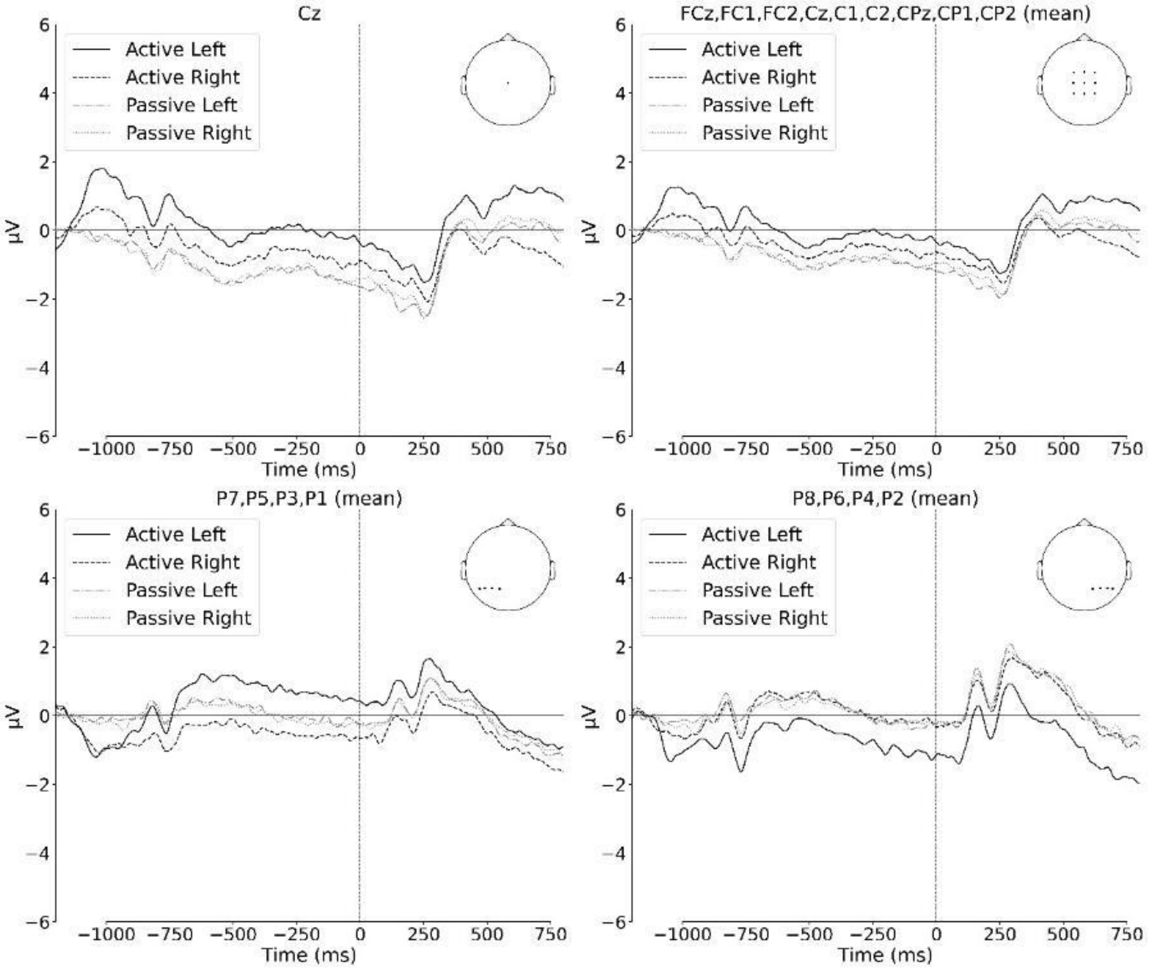


***Figure 20:*** *Grand-average ERPs across different electrodes. The baseline period was defined between -1200 ms and -1100 ms, relative to the motor action occurring at -1000 ms. Time 0 marks the onset of Grating 1. 'Left' and 'Right' indicate the orientation of Grating 1. Active conditions correspond to trials in which participants pressed a button to generate the grating orientation (left or right-oriented), while Passive conditions correspond to trials in which the orientation was pseudo-randomly presented.*

Table 1

| Participant | Error indices dropped from behavioural data | Interpolated electrodes | | IC component excluded and component score | % epochs removed | N epochs dropped | Epochs manually dropped |
|---|---|---|---|---|---|---|---|
| P001 | 127, 134, 274 | None | 0 | IC000 ; 0.91 | 7,68% | 43 | 0, 37, 72, 85, 92, 95, 100, 116, 142, 144, 146, 153, 175, 221, 231, 239, 256, 282, 285, 287, 321, 363, 370, 383, 384, 401, 415, 432, 444, 445, 458, 459, 470, 481, 490, 498, 512, 518, 520, 521, 522, 541, 542 |
| P002 | 132 | None | 0 | IC001 ; 0.89<br>IC002 ; 0.15 | 0,89% | 5 | 37, 43, 295, 315, 393 |
| P003 | 90, 225, 226, 424 | None | 0 | IC000 ; 0.78<br>IC008 ; 0.29 | 13,39% | 75 | 2, 5, 7, 10, 36, 39, 42, 84, 85, 86, 87, 92, 95, 96, 98, 104, 106, 112, 115, 117, 119, 129, 130, 134, 140, 143, 154, 155, 158, 167, 191, 192, 207, 209, 217, 218, 219, 220, 221, 222, 223, 235, 236, 240, 246, 248, 250, 271, 274, 284, 292, 301, 313, 319, 344, 345, 347, 362, 399, 405, 406, 414, 426, 432, 464, 466, 478, 482, 495, 499, 500, 514, 543, 551, 554 |
| P004 | 112 | FT9;FT10 | 2 | IC000 ; 0.94<br>IC001 ; 0.25 | 1,07% | 6 | 317, 340, 341, 366, 539, 548 |
| P005 | None | O1;PO7;PO3 | 3 | IC000 ; 0.94<br>IC001 (saccades)<br>IC002 ; 0.35 | 0,71% | 4 | 132, 213, 275, 373 |
| P006 | 75 | TP10 | 1 | IC000 ; 0.54<br>IC001 (saccades) | 1,79% | 10 | 10, 36, 68, 157, 178, 386, 414, 439, 448, 479 |
| P007 | 43 | None | 0 | IC000 ; 0.66<br>IC001 ; 0.51<br>IC004 ; 0.30 | 5,36% | 30 | 10, 42, 55, 99, 108, 138, 153, 205, 213, 243, 270, 347, 375, 383, 390, 397, 404, 415, 418, 423, 431, 435, 462, 468, 470, 478, 511, 512, 525, 547 |
| P008 | 14, 269, 304 | None | 0 | IC002; 0.42<br>IC003; 0.1<br>IC004 ; 0.23 | 9,11% | 51 | 4, 14, 41, 56, 63, 107, 110, 155, 156, 157, 158, 159, 161, 179, 185, 190, 191, 199, 207, 216, 225, 271, 291, 293, 295, 296, 302, 303, 304, 305, 307, 308, 311, 312, 317, 319, 322, 332, 333, 338, 353, 372, 387, 407, 418, 427, 448, 468, 498, 507, 517 |
| P009 | None | None | 0 | IC000; 0.95<br>IC003 (saccades) | 0,36% | 2 | 107, 278 |
| P010 | 112 | None | 0 | IC000; 0.75<br>IC003 (saccades)<br>IC007 (saccades) | 1,07% | 6 | 7, 159, 218, 242, 316, 465 |
| P011 | 343, 552 | C6 | 1 | IC000; 0.81<br>IC002; 0.25 | 1,79% | 10 | 13, 15, 29, 38, 82, 109, 175, 222, 377, 556 |
| P012 | None | None | 0 | IC000; 0.85<br>IC006 (saccades) | 1,96% | 11 | 2, 61, 62, 72, 301, 361, 372, 382, 534, 535, 558 |
| P013 | 132 | None | 0 | IC000; 0.52<br>IC005 (saccades) | 1,25% | 7 | 90, 92, 288, 387, 392, 396, 486 |
| P014 | None | None | 0 | IC000; 0.63<br>IC001 (saccades)<br>IC007 (saccades)<br>IC008 (saccades) | 2,32% | 13 | 46, 96, 102, 105, 110, 196, 211, 292, 420, 448, 476, 504, 524 |
| P015 | None | None | 0 | IC000; 0.92<br>IC001 (saccades) | 1,25% | 7 | 28, 85, 230, 233, 346, 404, 478 |

| | | | | | | | | |
|---|---|---|---|---|---|---|---|---|
| P016 | 491 | None | 0 | IC000; 0.60<br>IC001 (saccades)<br>IC002 (saccades) | 7,14% | | 40 | 2, 14, 27, 67, 76, 85, 112, 113, 114, 116, 120, 136, 150, 155, 157, 159, 173, 175, 179, 195, 199, 273, 277, 303, 308, 314, 331, 336, 341, 344, 419, 483, 497, 499, 501, 515, 518, 531, 537, 559 |
| P017 | None | None | 0 | IC000; 0.75<br>IC001; 0.29 | 2,32% | | 13 | 85, 94, 140, 185, 241, 250, 256, 257, 281, 357, 364, 441, 487 |
| P018 | 95, 136, 161, 219, 3 | None | 0 | IC000; 0.73<br>IC001; 0.46<br>IC002 (saccades) | 3,57% | | 20 | 69, 217, 241, 252, 307, 310, 335, 346, 429, 432, 439, 469, 470, 471, 485, 486, 499, 509, 524, 538 |
| P019 | None | PO8 | 1 | IC001; 0.74<br>IC003 (saccades)<br>IC005 (saccades) | 0,89% | | 5 | 63, 67, 342, 502, 539 |
| P020 | 103, 531 | TP9;O1 | 2 | IC005; 0.81<br>IC009 (saccades) | 7,14% | | 40 | 0, 23, 24, 25, 27, 57, 89, 98, 140, 141, 155, 156, 157, 203, 205, 206, 216, 229, 232, 250, 251, 277, 289, 301, 305, 317, 330, 338, 362, 363, 372, 378, 379, 395, 416, 440, 460, 510, 524, 532, 558 |
| P021 | None | None | 0 | IC000; 0.90<br>IC001 (saccades)<br>IC018; 0.10 | 0,36% | | 2 | 8, 422, |
| P022 | 518 | None | 0 | IC000; 0.64<br>IC001 (saccades)<br>IC003; 0.09<br>IC013; 0.08 | 4,82% | | 27 | 20, 26, 31, 33, 34, 55, 79, 169, 255, 293, 294, 300, 309, 329, 364, 391, 393, 397, 409, 410, 418, 448, 449, 458, 522, 549, 553 |
| P023 | 30, 31, 32, 36, 144, 257, 446, 550 | None | 0 | IC000; 0.70<br>IC001; 0.39<br>IC003 (saccades) | 1,07% | | 6 | 1, 3, 4, 36, 52, 138 |
| P024 | None | None | 0 | IC001<br>IC004 (saccades)<br>IC012 (saccades) | 5,18% | | 29 | 5, 9, 11, 16, 19, 32, 41, 43, 47, 51, 57, 74, 78, 85, 91, 109, 111, 117, 125, 148, 150, 157, 250, 258, 375, 422, 437, 457, 532 |
